# MechanoMaST – a multimodal pipeline for spatially registering mechanical and transcriptomic tissue data

**DOI:** 10.64898/2026.08.29.747727

**Authors:** Linda Decker, Dmitrii Olisov, Nikolai Schleußner, Hendrik Wiethoff, Thomas Schmidt, Henrik Nienhüser, Thomas Moritz Pausch, Jan Oliver Korbel, Alba Diz-Muñoz

## Abstract

Spatial-omics workflows enable spatially-resolved molecular analysis within tissues. Despite the prognostic value of tissue stiffness, these approaches have not yet incorporated direct, absolute mechanical measurements. This omission reflects several challenges, including sample requirements, low throughput, and the need for specialized equipment and complex data registration.

Here, we introduce MechanoMaST (mechanics mapped to spatial transcriptomics), a workflow that combines absolute mechanical measurements with spatial-omics. It pairs atomic force microscopy-based nanoindentation stiffness maps with spatial transcriptomics maps from adjacent tissue cryosections. The two modalities are computationally co-registered to enable spatial correlation at 100 µm resolution with a sub-10 µm mean accuracy, providing ground-truth mechanical data directly linked to spatial gene expression.

We demonstrate MechanoMaST in human colorectal cancer liver metastasis, generating a spatial resource from 10 patients and revealing a five-gene stiffness signature. MechanoMaST is readily adaptable to other tissues across development and disease, and extendable to additional spatial-omics modalities.

## Introduction

Tissue stiffness is an active regulator of cell and tissue behaviour throughout development, homeostasis, and disease. For example, during development, stiffness gradients guide neural crest cell migration^1,2^, mesoderm invagination during gastrulation^3^, and stem cell lineage commitment^4^. Beyond its physiological roles, altered stiffness is both a consequence and a driver of disease progression, as well as a valuable prognostic tool. Increased tissue stiffness is a hallmark of cancer and is broadly unfavourable, promoting malignant processes including cancer cell proliferation, metastasis, drug resistance, as well as immune evasion, ultimately leading to tumour progression^5^. In organ fibrosis, increased stiffness causes the recruitment and activation of fibroblasts, which further progresses the pathology^6,7^. In contrast, in the brain, neurodegenerative pathologies such as Alzheimer’s disease are marked by reduced stiffness, and softening of the medial temporal lobe predicts cognitive decline^8,9^. Thus, a deeper understanding of tissue stiffness will not only allow us to better understand fundamental tissue biology, but also unlock the potential of tissue mechanics for prognostic and therapeutic approaches. Doing so requires tools that place tissue stiffness in a larger molecular context.

Multi-omics approaches, combining genomics, transcriptomics, proteomics, and other techniques, have become powerful tools for understanding biological systems and developing therapies for various diseases, including cancer^10–16^. With the rise of spatial multi-omics, this holistic view of cell state has extended to the cellular environment as well. However, these approaches have not yet been expanded to include mechanical properties, and no workflow currently provides direct, absolute stiffness measurements co-registered with spatial-omics data. This gap arises from three distinct challenges. First, mechanical measurements by atomic force microscopy (AFM)-based nanoindentation are inherently low-throughput and require specialized equipment and extensive user training, regardless of whether they are later linked to omics data. Second, combining mechanics with other spatial-omics modalities introduces a sample-compatibility conflict: AFM requires tissue to be measured fresh, whereas spatial transcriptomics (ST) and proteomics are typically performed on fixed tissue^17,18^. Third, even once both datasets are acquired, no established method exists to reliably register stiffness measurements to spatially resolved omics data at matching resolution, nor to quantify the resulting registration error — despite this being essential for distinguishing reliably mapped measurements from spurious ones. In the absence of a workflow that addresses these three challenges others have pushed the field forward with innovative strategies that estimate mechanical properties computationally rather than measuring them directly, or that perform registration only partially. One such study estimated mechanical properties (namely tension, pressure and stress tensors) via image-based mechanical force inference, an approach inherently limited to cell-rich areas, in a previously published seqFISH dataset of fixed mouse embryos^19^. Another study estimated the strain within fractured mouse bones via a micro-finite element model based on *in vivo* computed tomography images, which they then visually aligned with *ex vivo* ST data^20^. Additionally, a neural network estimated breast cancer stiffness from collagen stains, though this approach has not yet been combined with ST^21^. Together, these studies illustrate both the field’s interest in incorporating mechanics into the spatial multi-omics space and the difficulty of doing direct measurement, with compatible sample handling, and quantified accuracy.

Here, we provide a workflow that achieves this integration: mechanics mapped to ST (MechanoMaST). Specifically, we use AFM-based nanoindentation, the current gold standard in the field^22^, to acquire spatial maps of absolute stiffness values from 20-µm-thick tissue cryosections. Directly adjacent cryosections undergo an untargeted, spatial barcode-based ST workflow (10x Genomics Visium). We map both datasets via an image registration pipeline with a sub-10 µm mean accuracy. By linking each stiffness measurement to an ST spot, we reveal novel associations between gene expression and stiffness.

We demonstrate MechanoMaST in the highly clinically relevant context of human colorectal cancer (CRC) liver metastasis (LM). Increased tissue stiffness in CRC and its metastases correlates with reduced disease-free survival. This is particularly relevant for LM, the most common manifestation of metastatic CRC and a major cause of CRC-related death^23–26^. This stiffness arises from the tissue’s architecture: metastasis comprise parenchyma, composed mainly of cancer cells, and stroma, the non-cancerous compartment consisting predominately of cancer-associated fibroblasts (CAFs) surrounded by extracellular matrix (ECM). During the progression of cancer, the ECM is heavily remodelled by CAFs through the secretion, degradation, and crosslinking of its components including collagens^27^. Due to its high ECM content, the stroma is the main determinant of tissue stiffness and can drive drug resistance and immune exclusion^27–32^. Given this central role, stromal stiffness is a promising medical target and the focus of this study.

Numerous clinical trials target ECM stiffness-associated proteins^31,33,34^, but lack mechanical measurements linking stiffness to progression. Broad CAF depletion can worsen outcomes^35,36^, whereas targeting specific subpopulations has shown greater promise^37,38^. This suggests effective therapies should target stiffness-promoting subpopulations specifically, underscoring the need for mechanical measurements and molecular understanding in treatment stratification.

By applying our MechanoMaST workflow to 11 CRC LM samples from 10 patients, we identified five genes associated with increased tissue stiffness: *TGFBI*, *TFF3*, *TFF1, PRAP1* and *PHGR1*, representing candidate biomarkers or therapeutic targets for further investigation. Notably, we found no positive correlation between stiffness and classic ECM components such as COL1A1, highlighting that at this late disease state, collagen gene expression levels are not a suitable predictor of tissue stiffness. This underscores the continued need for direct mechanical measurements. Furthermore, our extensive dataset provides a resource for future studies of stiffness-associated gene programs in cancer.

Beyond this specific application, MechanoMaST can be applied to any tissue that sufficiently retains its mechanical properties upon freezing. Tissue mechanical properties are altered in many diseases, including fibrosis, cancer, and neurological disorders^7,9,23,27,31,39,40^. Extending MechanoMaST to other tissue types could therefore improve our understanding of these pathologies, and ultimately their treatability. Furthermore, the MechanoMaST pipeline provides a multimodal framework that can be extended by additional methods in adjacent sections. Possible future applications include proteomics, high-resolution ST, metabolomics and lipidomics, enabling a more comprehensive understanding of tissue mechanics in health, development, ageing and disease.

## Results

### A multimodal pipeline for spatially registering mechanical and transcriptomic tissue data

To gain a deeper, spatially informed understanding of the relation between tissue stiffness and gene expression levels, we developed MechanoMaST. We perform ST using the Visium platform, as it offers total mRNA coverage and is therefore not biased towards a pre-selected set of genes. For mechanical measurements, we chose AFM, as it allows the detection of absolute stiffness values in the form of the Young’s modulus: the ratio of stress to strain in elastic materials. However, these two techniques impose conflicting sample requirements: AFM maps require the tissue to remain unfixed in liquid for hours, whereas Visium requires the tissue to be fixed immediately after thawing to preserve RNA integrity. To resolve this, we combine AFM-based nanoindentation with sequencing-based ST in adjacent serial tissue cryosections rather than the same section (**Fig. 1a**). The acquired datasets are then spatially registered via a custom image registration pipeline based on landmarks (**Fig. 1b–f**), enabling the detection of associations between gene expression and tissue stiffness.

**Figure 1:**
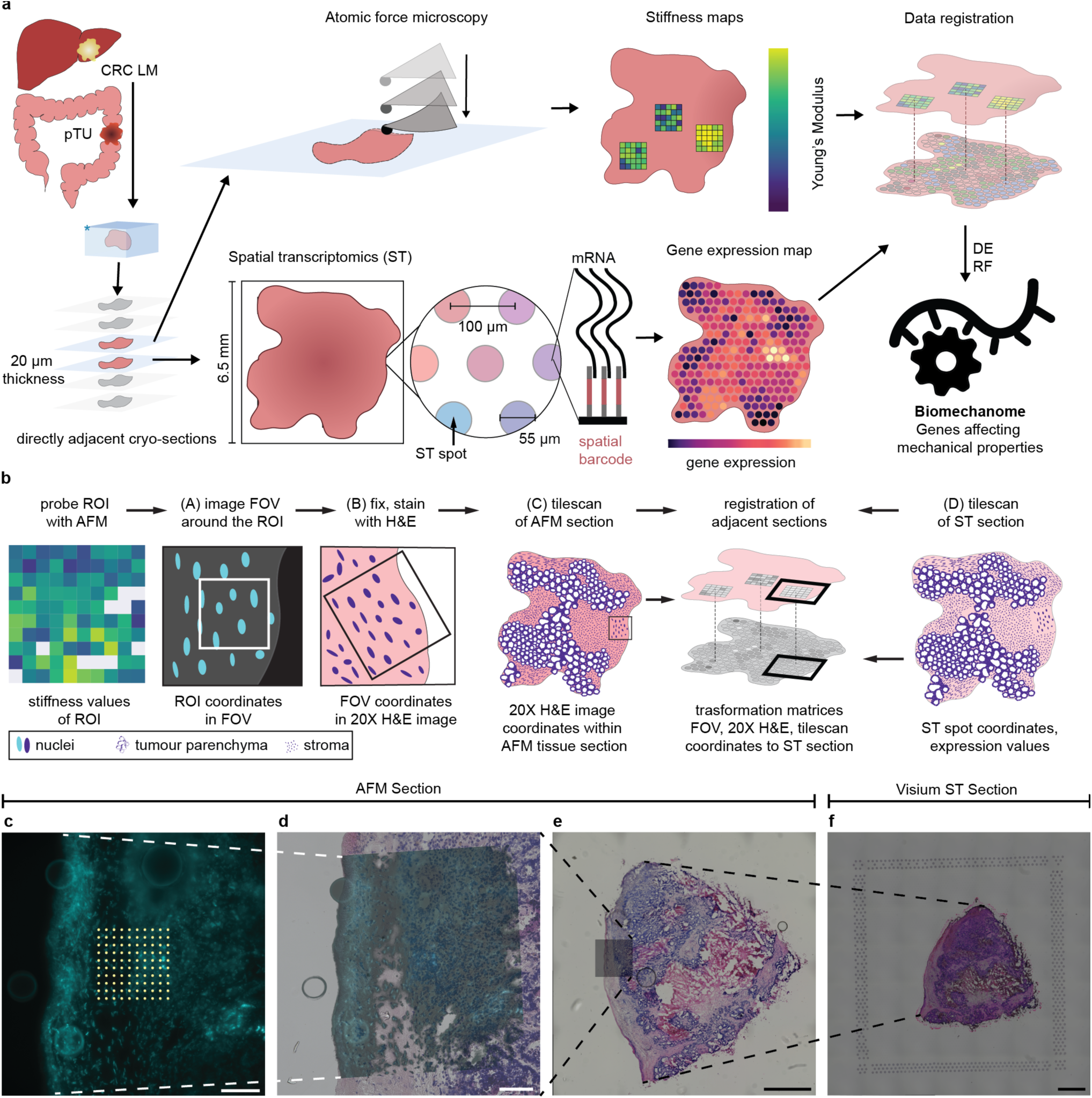
Overview of the MechanoMaST workflow and image registration pipeline. **a**, Schematic of the MechanoMaST workflow. CRC LM samples are freshly frozen in optimal cutting temperature compound on dry ice and cut into 20 µm thick sections. One of two adjacent cryosections is probed via AFM, yielding stiffness maps. The other section undergoes a sequencing-based ST workflow via the Visium platform, in which the cryosection is placed on a barcoded chip such that both mRNA sequences and spatial barcodes are captured. Stiffness and expression maps are then registered via the image registration workflow described in panels b–f. Genes associated with mechanical properties of tissue — the biomechanome — are identified via differential expression (DE) analysis and a random forest (RF) model, as described in Fig. 5. **b**, Schematic of the MechanoMaST image registration pipeline. Cartoon images A–D correspond to the representative images shown in panels c–f, respectively. **c–f**, Representative example of the registration pipeline for Patient 4. **c**, Unfixed, Hoechst-stained field of view (FOV) during AFM measurements; white dots indicate measurement coordinates. **d**, The image in c, affine-transformed and overlaid onto an image of the same FOV after H&E-staining. **e**, The image in d, scaled by a factor of 0.5 and template-matched onto a methanol-fixed, H&E-stained tile-scan of the same tissue section. **f**, Methanol-fixed, H&E-stained tile-scan of the section adjacent to c–e, placed on a barcoded chip with a fiducial frame as part of the Visium ST workflow; the image in e is mapped onto it via affine transformation. LM: liver metastasis; pTU: primary tumour; FOV: field of view. White scale bars: 100 µm; black scale bars: 1 mm.

In Visium, spatial information is encoded by barcodes organized into spots of 55 µm diameter, spaced 100 µm apart (centre-to-centre). An image of the tissue section is also acquired, to which the ST data is subsequently mapped. AFM measurements, by contrast, probe a much smaller area: to average stiffness over a wider tissue region per indentation, we equipped cantilevers with 25 µm silica beads, which, at an indentation depth of 250 nm, results in an estimated contact radius of 2.5 µm with the sample. AFM also enables simultaneous fluorescence imaging of the sample, allowing the acquired stiffness data to be mapped to images in the same way as Visium. Given this difference in scale, between AFM’s highly local probing and Visium’s comparatively large capture area, we set our AFM measurement spacing relative to the Visium resolution, placing indentations 20 µm apart in a grid-like manner. This reduces measurement time per region of interest (ROI) while ensuring multiple AFM measurements are mapped to each ST spot, averaging out local stiffness variations.

Since both modalities can be readily mapped to corresponding images, ST and AFM data are aligned through an image registration pipeline (**Fig. 1b**). In the case of the ST data, the mapping to an image, (here a widefield image of a Hematoxylin and Eosin (H&E)-staining), can be automated via the SpaceRanger pipeline (10X Genomics), which detects the fiducial frame of the barcoded chip on which the H&E-stained tissue is imaged before undergoing the RNA extraction and sequencing pipeline (see **Methods** for details). Based on the corner positions of the fiducial frame, the ST spots are mapped onto the H&E image.

To map AFM data to images, we stain the unfixed tissue with Hoechst, a nuclear marker, and acquire epifluorescence images of the probed regions before performing the AFM measurements. We centre the cantilever tip on the imaged field of view (FOV) before probing, so that each AFM measurement’s position within the grid directly gives its pixel coordinates in the image. To bridge the gap between the unfixed Hoechst image showing the AFM-probed FOV and the H&E-stained tile-scan of the adjacent tissue section, we fix and H&E-stain the AFM section after the measurements. AFM sections are thus imaged three times in total: unfixed and Hoechst-stained FOV during AFM measurements (**Fig. 1c**); fixed and H&E-stained FOV (**Fig. 1d**); fixed and H&E-stained tile-scan (**Fig. 1e**). The ST section is imaged only once as a fixed and H&E-stained tile-scan on the fiducial frame (**Fig. 1f**). These four images are then registered step-wise via affine transformations and template matching (see **Methods** for details). The resulting transformation matrix can then be applied to convert the original AFM measurement coordinates into Visium space, producing a dataset that links stiffness values to spatial gene expression.

### MechanoMaST reliably registers mechanical and transcriptomic data across adjacent CRC LM sections

We chose to demonstrate MechanoMaST in CRC LM for two reasons. First, CRC LM stand at the centre of significant clinical need^26^, and we have shown that reducing tissue stiffness is a promising approach for increasing patient survival: patients with CRC LM who took renin–angiotensin system inhibitors for hypertension had softer tumours, responded better to treatment with the VEGF antibody bevacizumab, and showed increased survival compared to normotensive patients or those on other antihypertensives^24^. Second, CRC LM cryosections are a well-suited technical model: we have previously shown that the mechanical properties of CRC LM stroma, as measured by AFM, are preserved in frozen tissue compared to matched fresh samples^24^.

As described above, MechanoMaST uses directly adjacent cryosections rather than the same section for AFM and ST acquisition. While previous work has reported a high Pearson correlation coefficient (0.98) for gene expression between adjacent mouse brain sections spaced 50 µm apart^41^, whether the same held true for tissue stiffness remained untested. We therefore performed AFM measurements on three adjacent sections and found that the stiffness pattern was conserved between them: differences between regions *within* the same section were larger than differences between the *same* region measured across different sections (**Fig. 2a-e**). Together with the small 20 µm section thickness relative to the 100 µm centre-to-centre distance of ST spots, this justified the use of adjacent tissue sections for the MechanoMaST workflow.

**Figure 2:**
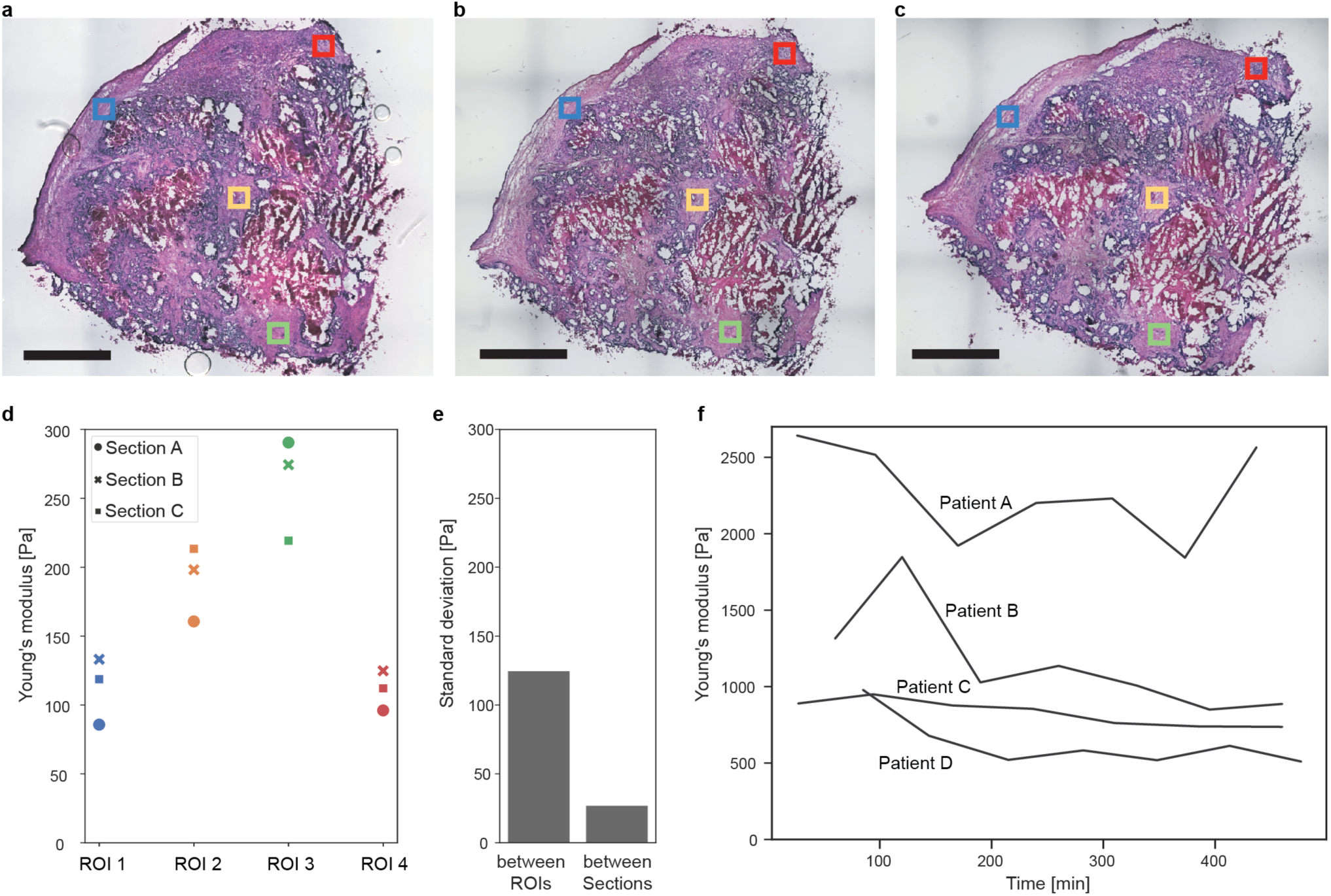
Tissue stiffness is stable across space and time. **a–c**, Directly adjacent 20 µm human CRC LM cryosections stained with H&E; coloured squares indicate the four ROIs probed via AFM. Scale bars: 1 mm. **d**, Average stiffness of each ROI across the three sections shown in a–c. A two way ANOVA with sections and ROIs as factors was performed: section: p = 0.4881, ROI: p = 0.0015 **e**, Standard deviation between ROIs and between sections shown in a–c and quantified in d. (n = 4 ROIs, 3 sections). **f**, Tissue stiffness over time (after thawing) in four patients (Patient A–D). One 200 × 200 µm ROI was probed every 20 µm, up to 7 times in a row per patient; each data point represents the average stiffness of the ROI at that time point. Only coordinates yielding a stiffness value at all time points were included. The sample shown in a–c and Patient D in panel f are from different tissue blocks of the same patient. Sections adjacent to those shown in a–c were processed with the MechanoMaST workflow as Patient 4; adjacent sections from Patient A and Patient B in panel f were processed as Patient 2 and Patient 1, respectively.

We further performed time-course experiments and observed only a moderate decrease in tissue stiffness over the course of the experiment (**Fig. 2f, Supplementary Fig. 1**).

### Quality control and error determination of the MechanoMaST workflow

We performed MechanoMaST on 11 CRC LM samples from 10 patients; from patient 7 samples from the centre (7a) and the periphery (7b) of the same metastasis were processed. Across samples, we observed high spatial variability in terms of morphology (**Fig. 3a-d**), gene expression (**Fig. 3e,f**; exemplified here by *COL1A2* and *FN1*, two common ECM genes), and stiffness (**Fig. 3g,h**). This intra-patient variability and spatial heterogeneity in stiffness and gene expression underscore the need for accurately mapped, spatially resolved data to capture the tissue’s complex organization (**Supplementary Fig. 2)**. Thus, to quantify the mapping accuracy of MechanoMaST, we performed an error propagation throughout the image registration steps (**Fig. 3i-n**, for details see **Methods**).

**Figure 3:**
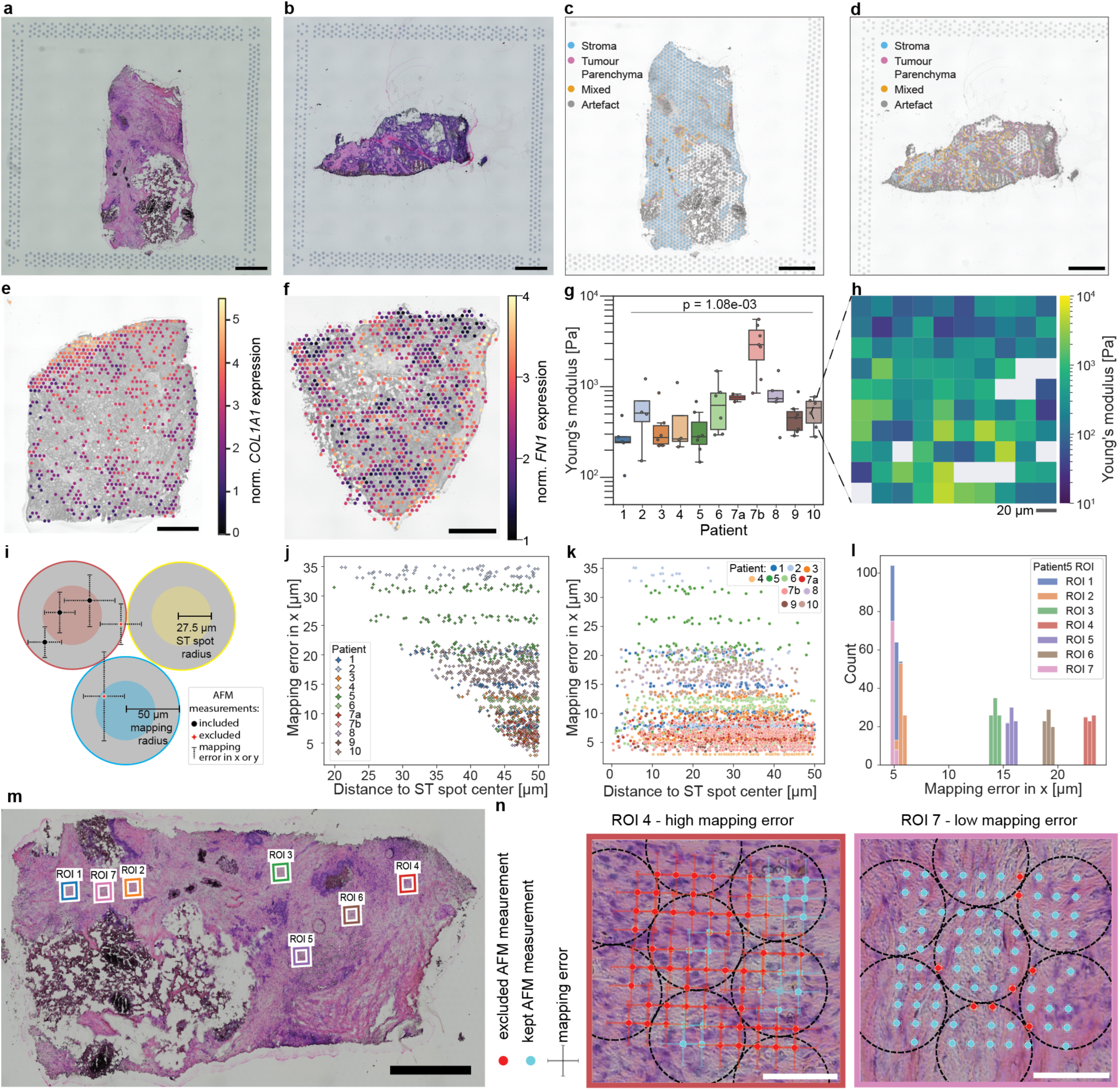
Spatial heterogeneity of CRC LM samples and MechanoMaST mapping accuracy. (n = 11 CRC LM samples from 10 patients; representative examples shown). **a,b**, CRC LM samples from Patient 5 (a) and Patient 8 (b) on Visium slides; methanol-fixed, H&E-stained. **c,d**, Pathologist’s morphological annotations of the samples shown in a,b. For annotation criteria see **Methods. e,f**, Normalized spatial expression of *COL1A2* (e) or *FN1* (f) in samples from Patient 2 and Patient 4, respectively; each dot represents one ST spot. **g**, Stiffness (Young’s modulus) across samples; each dot represents the average stiffness of one ROI (2–7 ROIs per sample, 16–162 measurements per ROI, most commonly 100; see **Supplementary Table 3** for exact counts per ROI). Samples differed significantly in stiffness (Kruskal–Wallis test; see **Methods** for details of test selection). **h**, Stiffness map acquired by AFM indentation; grey areas indicate measurements where the force-distance curve could not be reliably fitted with a Hertz model. Measurements are spaced 20 µm apart. **i**, Schematic of the mapping error exclusion criterion: AFM measurements that remain assigned to the same ST spot after being shifted by their mapping error are considered reliably mapped and retained; measurements reassigned to a neighbouring spot under this shift are excluded. **j,k**, Mapping error in x for excluded (j) and retained (k) AFM measurements, as a function of distance to the ST spot centre. **l**, Mapping error in x for Patient 5 (n = 556 measurements across 7 ROIs). **m**, All AFM-probed ROIs for Patient 5; mapping was more accurate toward the left side of the sample (see **Supplementary Fig. 3** for details). **n**, Close-ups of ROI 4 and ROI 7, showing mapped AFM measurement coordinates and their corresponding mapping errors (half the standard deviation derived from error propagation; see **Methods**). Black scale bars: 1 mm; white scale bars: 100 µm.

Quality control parameters for the individual techniques AFM and ST and a summary of patient characteristics can be found in **Supplementary Tables 1-3**. In short, we acquired 16675 ST spots from 11 tissues, with a median of 955 genes per spot per sample. 785 spots were excluded due to a low read number or a high percentage of detected mitochondrial genes, leaving 15890 spots. From our AFM measurements, 4584 out of 5810 recorded force-distance curves could be fitted with a Hertz model, with an average fit root mean square (RMS) residual of 14.67 pN. Since we specifically probed the stroma, we curated a set of 297 pathologist-annotated, AFM-probed stroma spots across all 11 samples for correlative analysis.

Because mRNA released between ST spots on a Visium slide may diffuse and be captured by a neighbouring spot, we also included AFM measurements mapped to the regions between spots in our analysis. Since neighbouring ST spots are spaced 100 µm apart (centre-to-centre), the midpoint between adjacent spot centres lies 50 µm from each centre. We therefore mapped every AFM measurement within 50 µm of an ST spot’s centre to that spot (**Fig. 3i**), extending beyond the spot’s 27.5 µm physical radius to maximise our mechanical data yield and statistical power. To ensure this procedure does not compromise our data quality, we thoroughly analysed our mapping accuracy. For each measurement, we propagated the errors from both affine transformation steps (**Supplementary Table 3**). Only those measurements that still mapped to the same ST spot when moving their spatial coordinates by their mapping error (half a standard deviation) were considered reliably mapped and kept for downstream analysis (**Fig. 3i-k**). Thus, for measurements that landed in the centre of an ST spot, a higher mapping error was tolerated (**Fig. 3i-k**). The extent of the mapping error depended largely on the probed ROI (**Fig. 3l, Supplementary Table 3**) and the position of the ROI within the tissue and with respect to the registration landmarks (**Fig. 3m-n**, **Supplementary Fig. 3**). Nevertheless, as described below smaller mapping radii produced similar results (**Supplementary Fig. 4**).

### MechanoMaST reveals that classic ECM components show limited correlation between gene expression and tissue stiffness

We next sought to quantify the relationship between gene expression and stiffness in CRC LM. As a starting point, we drew on prior literature to select relevant ECM components. A mass spectrometry-based comparison of human CRC LM and healthy liver tissue identified the following ECM-associated peptides as most upregulated in CRC LM in descending order: COL1, COL1A2, COL3A1, FBN1, FN1, COL12A1 and COL5A1^28^. We therefore focused first on the genes encoding these seven proteins, terming their combined expression level the ECM score (**Fig. 4a**).

**Figure 4:**
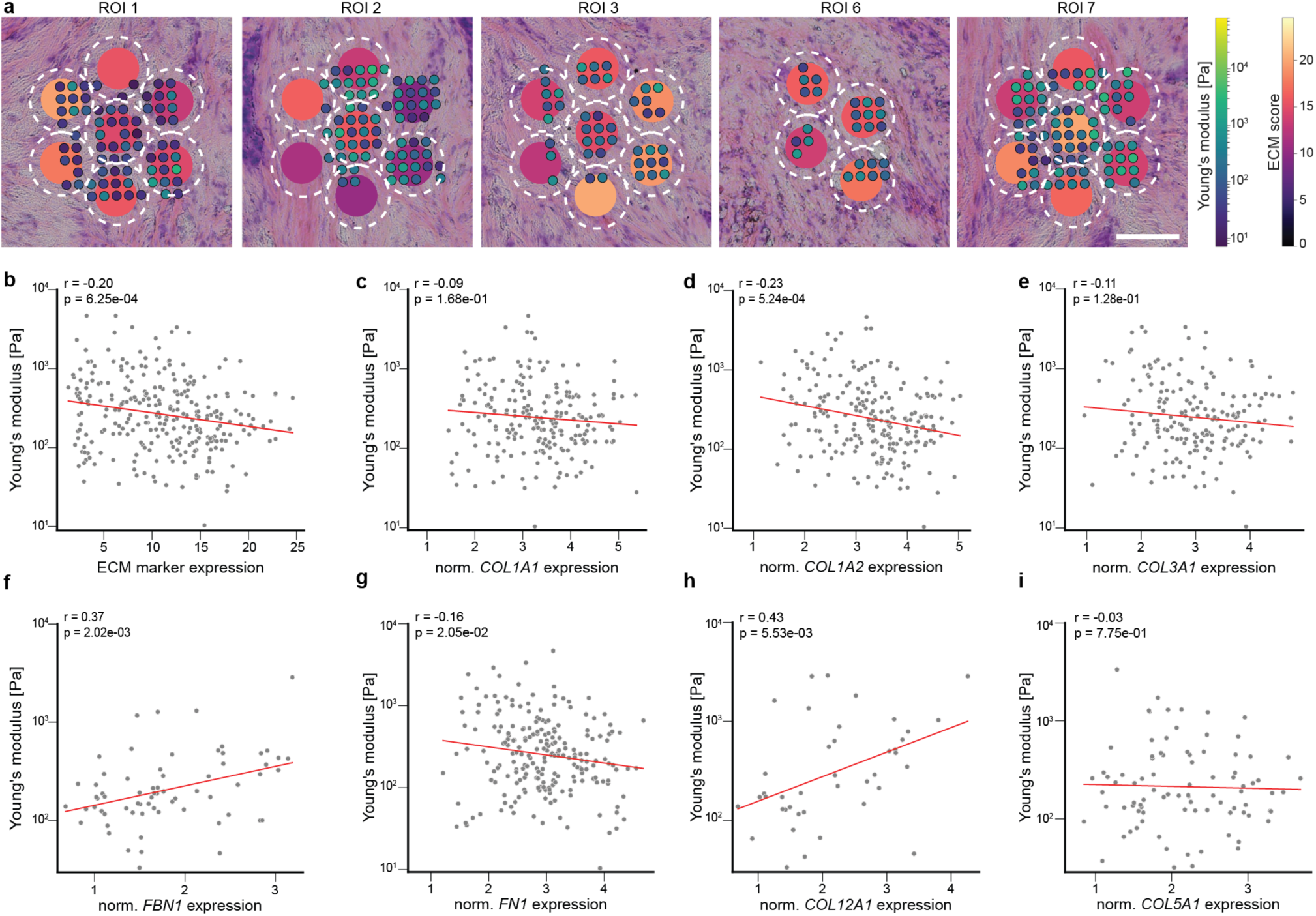
Classic ECM components show limited correlation between gene expression and tissue stiffness. (n = 297 ST spots). **a**, ECM score and AFM-probed stiffness mapped onto an H&E-stained CRC LM sample; representative ROIs from Patient 5 are shown. Large spots represent ST spots; small spots represent AFM measurements. Dashed white circles indicate which ST spot each AFM measurement is linked to. Spots with only one mapped measurement were excluded from downstream analysis. The ECM score is the summed lognorm expression of *COL1A1*, *COL1A2*, *COL3A1*, *FBN1*, *FN1*, *COL12A1* and *COL5A1*. **b–i**, Linear regression of Young’s modulus against the lognorm expression of the ECM score (b) or individual ECM score genes (c–i); each dot represents one ST spot. Only spots with non-zero expression of the respective genes are shown. ECM markers: 279 spots, *COL1A1*: 222 spots, *COL1A2*: 226 spots, *COL3A1*: 189 spots, *FBN1*: 67 spots, *FN1:* 218 spots, *COL12A1*: 41spots and *COL5A1*: 89 spots. r: Pearson’s coefficient. Scale bar: 100 µm.

To assess how individual genes contributing to the ECM score relate to stiffness, we averaged the stiffness of all AFM measurements mapped to the same ST spot and performed linear regressions (**Fig. 4b-i**). There was a significant negative correlation between the overall ECM score and stiffness (**Fig. 4b**). However, the small Pearson coefficient for this correlation suggests it is unlikely to be biologically meaningful. Surprisingly, we found no correlation between *COL1A1, COL3A1,* and *COL5A1* gene expression and stiffness (**Fig. 4c,e,i**) and a negative one for *COL1A2 and FN1* (**Fig. 4d,g**). Only *FBN1* and *COL12A1* showed the expected significant positive stiffness correlation (**Fig. 4f,h**).

While ECM signatures have often been used to make claims about tissue stiffness ^38,42,43^, we show that classic structural ECM components are not the primary drivers of stiffness at the RNA level. Other factors, potentially ECM regulators, interaction partners, or modifiers, may instead underlie this association. This highlights the importance of incorporating direct mechanical measurements into workflows. Furthermore, it underscores the value of datasets such as the one presented here to identify genes that stratify tissue stiffness.

### DE and RF analysis reveal *TGFBI*, *TFF3*, *TFF1*, *PRAP1* and PHGR1 as stiffness-linked genes in CRC LM

Given the inconsistent and counterintuitive correlations between the most abundant ECM components and tissue stiffness, we decided to adopt completely unbiased approaches to identify genes that are linked with malignant tissue stiffening in cancer. To identify genes associated with high stiffness, we applied two complementary approaches (differential expression (DE) analysis and a random forest (RF) model). For the DE analysis, we selected three different thresholds (**Fig. 5a**) for what is considered a stiff spot: 260 Pa (**Fig. 5b**), 520 Pa (**Fig. 5c**), and 780 Pa (**Fig. 5d**). We chose 260 Pa as an initial threshold as it distributed a similar number of spots into the soft and stiff categories, and then two and three times that value, 520 Pa and 780 Pa, respectively, as additional thresholds to minimize threshold-induced biases in the analysis. Applying the differential expression analysis using smaller mapping radii produced similar results (**Supplementary Fig. 4**).

**Figure 5:**
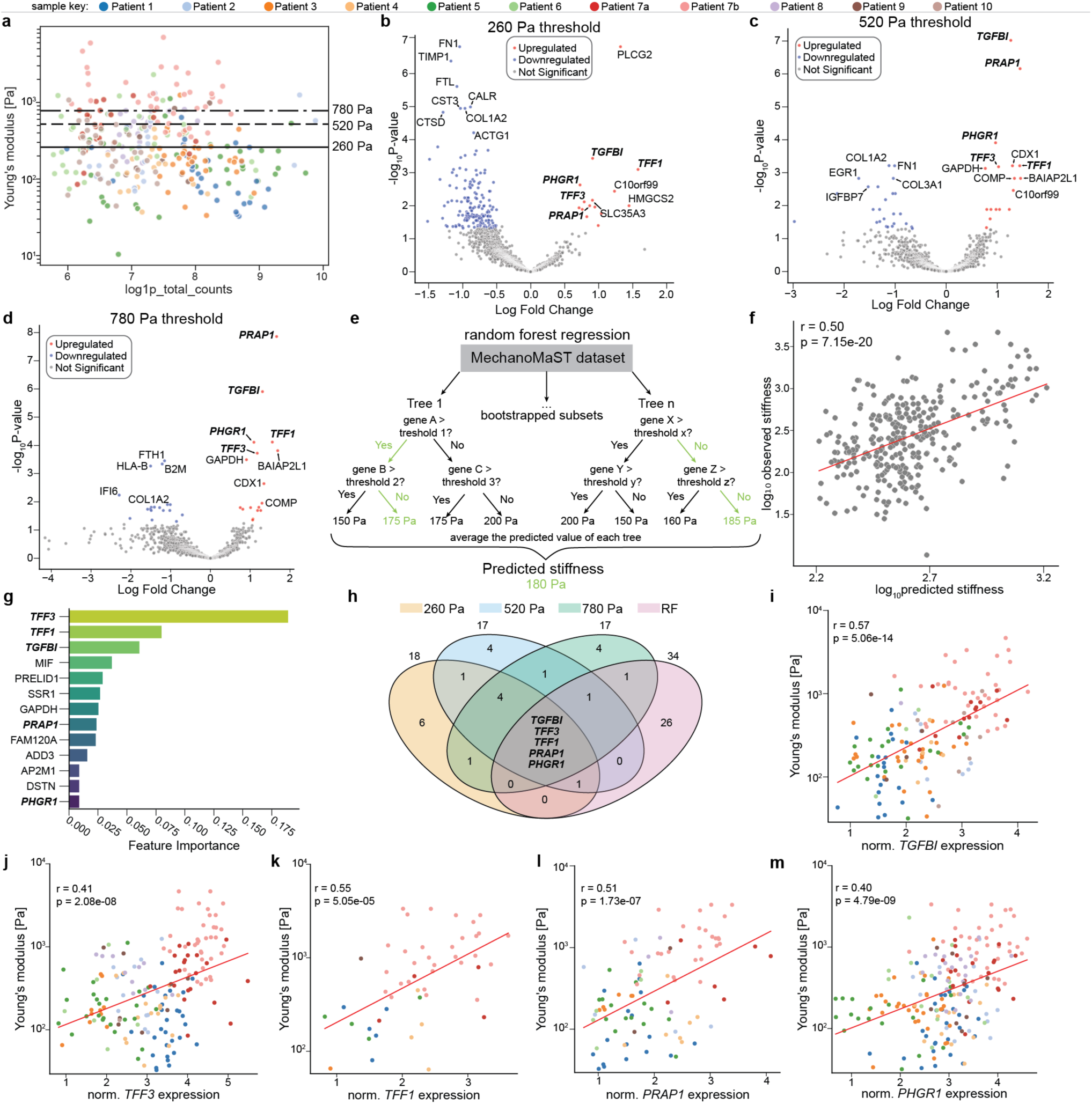
Identification of *TGFBI, TFF3, TFF1, PRAP1 and PHGR1* via differential gene expression (DE) analysis and a random forest (RF) model. (n = 11 CRC LM samples from 10 patients; 297 stroma spots probed by AFM). **a**, Overview of all stiffness-probed ST spots of the MechanoMaST dataset after filtering; horizontal lines indicate the three thresholds used for differential expression analysis. **b–d**, Differential expression analysis between soft and stiff spots at each threshold (b, 260 Pa; c, 520 Pa; d, 780 Pa); each dot represents one gene. Red dots refer to genes upregulated in stiff spots and blue dots refer to genes downregulated in stiff spots. **e**, Principle of the RF model. **f**, Correlation of observed and RF-predicted stiffness; each dot represents one ST spot, n = 297 spots. **g**, Top 13 genes by feature importance in the RF model; higher feature importance indicates greater contribution to the stiffness prediction. **h**, Number of target genes identified by each approach (DE and RF), and the five genes common to all approaches. **i–m**, Correlation of lognorm gene expression and measured stiffness for *TGFBI* (i, 144 spots), *TFF3* (j, 176 spots), *TFF1* (k, 50 spots), *PRAP1* (l, 93 spots) and *PHGR1* (m, 203 spots); each dot represents one ST spot, only spots with non-zero expression are shown. r: Pearson’s coefficient.

Additionally, we trained a random forest model on our dataset to predict stiffness based on gene expression (**Fig. 5e-g, Supplementary Fig. 5**). The model uses bootstrapping to create subsets of the dataset and builds decision trees. After the model is trained, it predicts stiffness by averaging the predictions of all decision trees (**Fig. 5e**). Extracting feature importance, which reflects how much each gene contributed to the stiffness prediction, allows the RF model to identify genes correlating with high stiffness, providing an alternative discovery approach (**Fig. 5g**). Complete gene lists for all approaches can be found in **Supplementary Tables 4-9.** The DE analysis and the RF model converged on the same five genes: *TGFBI, TFF3, TFF1, PRAP1* and *PHGR1*, a striking consensus across independent approaches (**Fig. 5h**).

Having identified five candidate genes, we verified that all of them had a significant positive correlation between gene expression and tissue stiffness (**Fig. 5i-l, Supplementary Fig. 5a-e**). Next, we compared the expression across patients (**Supplementary Fig. 6**), and found that TFF1 is predominantly expressed by Patient 7, while the other genes were highly expressed by multiple patients. This suggests a patient-specific effect for *TFF1*, whereas *TGFBI, TFF3, PRAP1* and *PHGR1* may contribute more broadly to tissue stiffening. To the best of our knowledge, only *TGFBI* has previously been implicated in malignant tumour stiffening^44^, while the other candidates are novel. These findings underscores the power of MechanoMaST as an unbiased discovery approach, capable of uncovering previously unrecognized stiffness-linked genes.

## Discussion

With MechanoMaST we introduce a pipeline that spatially co-maps direct mechanical measurements with transcriptomic landscapes, expanding the scope of spatial multi-omics with spatial mechanomics. Unlike prior approaches that infer mechanical properties computationally or align datasets only visually, MechanoMaST provides absolute stiffness measurements co-registered with spatial gene expression at 100 µm resolution with sub-10 µm accuracty.

MechanoMaST relies on measurements in adjacent cryosections. We show that tissue stiffness is sufficiently well preserved across directly adjacent CRC LM sections to justify acquiring ST and AFM data separately in this manner. Using consecutive cryosections for the AFM and ST workflows overcame the challenge of differing sample condition requirements. However, it created a need for accurate spatial registration. The use of directly adjacent tissue sections, including their registration via affine transformation, is common in the spatial multi-omics field^45–47^. However, in our case, the AFM and Visium sections were first imaged in completely different conditions, thus requiring a multi-step custom registration pipeline. Moreover, unlike previous approaches that align mechanical and omics data through visual inspection alone, we explicitly quantified mapping accuracy through error propagation, removing 23.2% of unreliable data points and leaving a refined, high-accuracy dataset.

Due to the complex 3D-structure of tumours, adjacent tissue sections can differ in their 2D morphology in a non-linear way, that affine transformations, limited to rotation, shearing, translation and scaling, cannot capture. Future work could address this by incorporating nonlinear deformation models into the MechanoMaST workflow, which should improve mapping accuracy^48^.

A general challenge of the workflow is the low throughput of AFM measurements: acquiring all stiffness maps took on average 6.5 hours of pure recording time per sample, allowing only a fraction of the sample to be probed. We placed the indentations 20 µm apart, resulting in mapped mechanics for a mean of 27 ST spots corresponding to 5–93% of stromal ST spots per sample. The spacing was deliberately set relative to the 100 µm centre-to-centre spot distance of Visium, so that multiple AFM measurements were averaged per spot, while maximising the probed stromal area, generating a balanced, unbiased resource for identifying stiffness-associated genes.

With the emergence of high-resolution ST approaches^49^, future applications with a higher-resolution version of MechanoMaST using more closely spaced AFM measurements would be feasible. However, higher resolution introduces its own challenges: finer registration accuracy would be required to reliably assign AFM measurements to smaller ST spots, and enough AFM measurements per spot would still be needed to average out measurement noise.

We demonstrated the MechanoMaST workflow on clinically relevant human CRC LM samples, generating a first resource for future research into tissue mechanics and disease. During the analysis of this dataset, we initially assessed how ECM-related genes, including collagens, correlated with tissue stiffness and observed predominantly negative or no correlations. This may seem surprising, given the positive correlation of ECM components and tissue stiffness previously observed at the protein level^21,50–52^. However, considering the exceptionally long half-life of ECM components such as human collagen which can persist in cartilage for decades^53^, the mRNA levels are not an appropriate proxy for ECM protein abundance^54–57^. For tumours, ECM protein lifetimes are unknown. Since our measurements capture tissue only at the time of surgery, ECM accumulation may have largely already occurred, with remodelling through cross-linking and degradation becoming more prominent than production and secretion. Consistent with this interpretation, others have observed a downregulation of ECM-related genes, including *COL3A1* and *FN1*, in breast cancer cells grown on a stiff hydrogel compared to on a soft one^58^. This hints that the observed negative correlation of ECM components with stiffness might not only be a consequence of the time of sampling but may itself be medically relevant. Here, our pipeline opens up the possibility to mechanistically study whether low expression of certain ECM genes marks completed ECM deposition. To further characterize the levels, spatial distribution, and crosslinking state of collagens and other ECM components, our workflow could be integrated with proteomic approaches, for example by subjecting an adjacent cryosection to spatial mass spectrometry, immunostaining, CNA35 labelling, or picrosirius red staining.

We then applied two independent analytical approaches (RF and DE) and various stiffness thresholds 260, 520 and 780 Pa). We robustly identified a five-gene signature (*TGFBI, TFF3, TFF1, PRAP1 and PHGR1*) consistently associated with high ECM stiffness in CRC LM.

*TGFBI* is our strongest candidate. Its corresponding protein, TGFBIp (also known as βig-h3), is part of the ECM and has been associated with thicker collagen fibres and increased tumour stiffness in pancreatic cancer mouse models^44^. It is tempting to speculate that TGFBIp controls ECM stiffness through collagen organisation or bundling. Furthermore, *TGFBI* is upregulated in CRC and negatively correlates with T-cell activation, CD8^+^ immune cell infiltration and survival rates^44,59^, suggesting that it contributes to an immune-exclusionary environment and would be a promising target for future cancer therapies.

This fits a broader pattern in the field: broadly depleting CAFs, for example via αSMA⁺ ablation or Shh knockouts, can worsen outcomes^35,36^, whereas targeting defined subpopulations, such as LRRC15⁺^37^ or YAP1⁺ CAFs (linked to ECM stiffness and poor survival in hepatocellular carcinoma)^38^, has shown greater promise. TGFBI represents exactly this kind of specific, mechanistically grounded target for future therapeutic development. This is also consistent with our own prior finding that reducing tissue stiffness itself improves outcomes in CRC LM: patients taking renin–angiotensin system inhibitors for hypertension have softer tumours, respond better to VEGF-antibody treatment (bevacizumab), and show increased survival rates compared to those on other or no antihypertensives^24^.

The other four candidates (*TFF3*, *TFF1, PRAP1* and *PHGR1*) have not been previously linked to tissue stiffness. TFF3 has, however, been associated at the protein level with higher cervical mucus plug viscoelasticity^60^. These genes therefore represent novel candidates whose potential mechanobiological roles require further investigation.

Whether our candidate genes are a cause or consequence of increased tissue stiffness remains an open question. To address this, MechanoMaST could be applied to animal tumour models at different disease stages, enabling the detection of genes whose upregulation precedes increases in tissue stiffness. Notably, *TGFBI*, *TFF3, TFF1* and *PRAP1* encode secreted proteins^61–63^, supporting a plausible mechanistic role in modulating stiffness through direct interaction with, or incorporation into, the ECM. Their extracellular localization further makes them accessible to targeting, for example via function-blocking antibodies, underscoring their therapeutic potential. This potential is exemplified by TGFBIp, where antibody-based targeting has already reduced tumour burden relative to controls in pancreatic cancer mouse models^44^. Expanding such experiments to CRC models, and to the remaining candidate genes, could identify new mechanomedicines capable of reducing both stiffness and tumour burden.

Remarkably, *TGFBI*, *TFF3* and *PRAP1* have also been associated with metabolic dysfunction-associated steatotic liver disease (MASLD), formerly known as non-alcoholic fatty liver disease^64–67^. MASLD is associated with both, increased liver stiffness and CRC LM progression^68,69^. Thus, our candidate genes might contribute to the metastatic potential of CRC and the increased tissue stiffness by promoting lipid and ECM accumulation in the liver. Extending MechanoMaST to patient-paired non-cancerous liver samples would offer a direct test of this hypothesis.

As MechanoMaST is unbiased and species-agnostic, we expect it to be broadly transferable to other tissues. Promising model systems include pancreatic adenocarcinoma, due to its exceptionally high stiffness, and fibrosis-related diseases with extensive ECM deposition, such as lung fibrosis, heart fibrosis, or MASLD. More broadly, any tissue from any species in health, disease, development or ageing that can maintain relevant mechanical properties throughout the freezing and measurement process can be subjected to MechanoMaST. A retention of the mechanical properties can be confirmed by performing comparative measurements in fresh and fresh-frozen samples as we have done before in CRC LM^24^. Individually, both AFM and Visium have already been applied to a myriad of tissues. Other sequencing or imaging based ST methods would also be compatible with our workflow. For Visium specifically, optimal permeabilization time can be readily determined using commercially available kits and manufacturer protocols. For AFM, parameters such as indentation depth, bead size, measurement speed, and time window need careful selection, a task well within reach for anyone experienced with the technique. To adapt MechanoMaST to other sample types, adjacent sections should first be probed with AFM to confirm that mechanical properties are conserved between them. If required, section thickness could also be further reduced.

Because MechanoMaST’s core workflow requires only two adjacent cryosections, additional modalities (such as proteomics, metabolomics, or epigenomics) can readily be incorporated, either through further adjacent sections or, where applicable, additional measurements on the same section. Here, we have shown that mechanical properties are conserved across at least 40-60 µm of tissue (three 20 µm sections), leaving at least one further section available for additional methods, or more if thinner sections are used. In doing so, MechanoMaST establishes tissue mechanics — the ‘mechanome’ — as a modality that can be spatially co-registered alongside the genome, transcriptome, proteome, and metabolome, completing a previously missing dimension of the spatial multi-omics landscape.

In this study, we show that direct mechanical measurements and spatial gene expression can be reliably co-registered across adjacent tissue sections to reveal biomechanical insights together with transcriptomic issue profiling. MechanoMaST is the first workflow to spatially map absolute stiffness values in Pascals (rather than relative or dimensionless mechanical estimates) to the local transcriptome (with registration accuracy quantified through error propagation rather than visual assessment), thereby addressing the long-standing lack of mechanical data in the spatial omics field. Our application of this workflow in CRC LM demonstrates its broad potential for both fundamental and medical research, enabling deeper insights into stiffness-associated genes and the identification of promising targets for mechanomedicines.

## ACKNOWLEDGEMENTS

We thank Sarah Katherine Foster for continuous feedback and critical discussions throughout the preparation of this manuscript. We thank Robert Prevedel, Anna Kreshuk, Vladimir Benes, Sinem Saka, Martin Bergert, Magdalena Glotzer, Roshni Banerjee and Markus Körbel for critical reading of the manuscript. We thank the EMBL Data Science Centre and GenTechDev OpenLab (especially Felix Schneider, funded by MULTI-SPACE, Health + Life Science Alliance Heidelberg Mannheim, and Sarah Kaspar) for guidance and support on error propagation and statistics. We additionally thank the EMBL Genomics Core Facility, the EMBL Advanced Light Microscopy Facility, and the EMBL Mechanical Workshop for support and advice. We are grateful for the tissue service at the General, Visceral, and Transplant Surgery, Heidelberg University Hospital, for transporting samples from the operating rooms to the hospital lab. We would also like to thank the patients and their families for donating tissue samples to this work.

## AUTHOR CONTRIBUTION

A.D.-M. and L. D. conceived the project. L.D. and A.D.-M. designed the experiments. L.D. performed all experiments, and developed and applied the image registration and data mapping pipelines. D.O. performed the analysis of the Visium data, differential gene expression analysis and built the random forest model. N.S. and L.D. freshly froze tissue samples. N.S., H.N., T.M.P., and T.S. coordinated patient consent and sample acquisition. H.W. performed the final ST spot annotation and provided expertise during the sample selection process. H.W., L.D., N.S, H.N., T.M.P. and A.D.-M. selected the samples for MechanoMaST together. L.D. and A.D-M. wrote the manuscript. All authors contributed to the interpretation of the data, read, edited and approved the final manuscript.

## Methods

### Patient samples

Samples were collected from patients who underwent curative resections of CRC LM at the Department of General, Visceral, and Transplant Surgery at the Heidelberg University Hospital between 2021 and 2024. All patients gave informed consent. This study was approved by the EMBL Bioethics Internal Advisory Committee (study number 2019-015) and the ethics committee of the Medical Faculty of the University of Heidelberg (study number S-708/2019). CRC LM were rinsed in PBS and embedded in optimal cutting temperature (OCT) compound (Sakura, 4583) in cryomolds (Sakura, 4566). The samples in cryomolds were immediately placed in a pre-cooled custom aluminium mould, engulfing the bottom and sides of the samples on dry ice. This freezing process avoids toxic isopentane baths and ensures a uniform contact area between the sample and the dry ice. Samples were transported on dry ice and stored at -80 °C.

### Cryo-sectioning

Human CRC LM samples were sectioned at a CM3050 S cryostat (Leica) using MX35 Ultra microtome blades (epredia, 3053835). The chamber temperature was kept at -20 °C and the object temperature at -10 °C. For MechanoMaST two directly adjacent sections (for patient 6, one section apart) were collected for AFM and ST. Samples intended for ST were scored if necessary, so that the sections fit into the capture area of the gene expression slides they were collected on (see spatial transcriptomics section). Sections for AFM measurements were placed on top of an 18 mm diameter aluminium cylinder and transferred into pre-coated glass-bottom dishes by very gently pressing the dish onto the sample. The day before sectioning, the dishes (wpi, FD35-100) were plasma-cleaned (GaLa Instrumente) air-based for 1 min and coated with 200 µl of 0.28 mg/ml Cell-TAK (Corning, 354240) at 37 °C overnight. Before tissue sectioning, the coated dishes were rinsed with Milli-Q water and let dry under a fume hood.

### Spatial transcriptomics (ST)

ST was performed on 11 CRC LM samples from 10 patients via the Visium platform (10X Genomics, 1000187, Standard Visium v1 for fresh-frozen tissue). According to the manufacturer’s recommendation, the permeabilisation time was optimised with the tissue optimisation kit (10X Genomics, 1000193) following the manufacturer’s guidelines (10X Genomics, CG000238 Rev E). In short, 20µm thick CRC LM cryosections were placed on tissue optimisation slides, stained with Hematoxylin and Eosin (H&E), imaged at a Nikon Ti2 microscope via a 10X tile scan and then lysed for different amounts of time. The mRNA binds to the slide via poly-A capture and is reverse-transcribed into fluorescent cDNA. The permeabilisation time leading to the highest fluorescence signal-to-noise ratio indicates the best mRNA yield and the optimum permeabilisation time. For our CRC LM samples, a permeabilisation time of 14 min was determined. For the H&E staining (10X Genomics, CG000160 Rev C), the isopropanol incubation step was reduced to 30 seconds.

The Visium ST workflow was performed per the manufacturer’s instructions (10X Genomics, CG000239 Rev F). In short, 20µm thick CRC LM cryosections with an RNA integrity number of at least 7 were placed on gene expression slides, stained with H&E (10X Genomics, CG000160 Rev C; isopropanol incubation reduced to 30 seconds), imaged without coverslips at an inverted Nikon Ti2 microscope with a Plan Apo λ 10X objective as a tile scan and permeabilised. The mRNA was captured on the gene expression slides and reverse-transcribed into cDNA, from which a sequencing library was prepared. The libraries were sequenced on Illumina NextSeq2000 machines in P3 flow cells with 50 bp paired-end sequencing, yielding between 56,000 and 138,000 mean reads per spot. The resulting FASTQ files were pre-processed together with the H&E images on the fiducial frame with SpaceRanger (10X Genomics, Version 2.1.0, available at https://www.10xgenomics.com/support/software/space-ranger/downloads). This software maps transcripts to ST spots using spatial barcodes and maps the spots to image coordinates using the fiducial frame as a reference.

For patient 4, the automatic orientation detection with SpaceRanger did not work. Therefore, the image of the Visium slide was manually rotated and flipped to match the orientation that SpaceRanger expects. For patients 4, 7 and 10 the corners of the fiducial frame were manually selected in Loupe Browser (v8.0.0, 10X Genomics) to refine the mapping.

Each ST spot was categorised by pathologist Dr. med. Hendrik Wiethoff as stroma, tumour parenchyma, mixed, healthy tissue, or artefact according to the following criteria:

- Stroma: > 70% of the spot is located within stroma
- Tumour Parenchyma: > 50% of the spot is located within tumour parenchyma
- Healthy: > 70% of the spot is located within healthy-appearing liver tissue
- Artefact: > 50% of the spot is located within a region with cryo-artefacts, necrosis, edema, or outside of the tissue
- Mixed: The spot does not fit the previous criteria

ST data were processed using Scanpy (v1.9.5) in Python (v3.9.18). Spots with fewer than 200 counts or more than 12,500 transcripts were removed. Only capture spots annotated as stroma by the pathologist were retained for further analysis. Additionally, genes expressed in less than 10% of the spots were filtered out. The raw count matrix was saved for differential expression analysis. For dimensionality reduction and visualisation, the count matrix was then normalized using the scanpy.pp.normalize_total function and log-transformed, then the UMAP algorithm was applied using the scanpy.tl.umap function. Clustering of cells within individual samples was done with the Leiden algorithm applied by scanpy.tl.leiden.

### Atomic force microscopy (AFM)

20 µm human CRC LM cryosections were thawed and stained with as low agitation as possible with Hoechst (Sigma, B2261; final concentration 2 µg/ml) in COPS (10X: cOmplete EDTA-free protease inhibitor cocktail (Roche, 11873580001; 1 tablet per 5 ml), 10% Penicillin/Streptomycin (Gibco, 15140-122) in PBS; store at -20 °C, thawed and diluted to 1X in PBS before usage) for 5 min. During measurements, samples were kept at 25 °C in 1 ml COPS.

AFM measurements were carried out at a CellHesion200 (Bruker) integrated into an Eclipse Ti inverted light microscope (Nikon). Cantilevers at position D of MLCT-O10 probes (Bruker) were calibrated via contact-based calibration in air, and their spring constants (0.024 – 0.042 N/m) were determined via thermal oscillation before attaching a 25 µm silica bead (microParticles, SiO2-R-25.0) with two-component glue (UHU Plus Schnellfest) at least 1 day before the measurements. Cantilevers in other positions were broken off using forceps. On the day of the measurement, the cantilever was coated with 1% Pluronic F-127 (Sigma, P2443) for 1 hour at 25 °C followed by a contact-based recalibration of the cantilever sensitivity in PBS. Stiffness measurements were conducted in the tumour stroma spaced 20 µm apart in a grid shape. For registration purposes, each probed area was imaged at 20X magnification (Nikon S Plan Fluor ELWD 20x/0.45 Infinity DIC N1 Objective) before the start of the measurements. Depending on the size of each particular stroma island, 16 – 400 indentation curves were acquired in each grid, yielding 255 – 700 force curves per sample. The samples were indented with a speed of 0.4 µm/s as previously described^70^ up to a set point of 5 nN.

Upon completion of the AFM measurements, the sections were fixed in -20 °C methanol for 30 min and stained with H&E (10X Genomics, CG000160 Rev C) with a reduced isopropanol incubation of 30 seconds. To adapt the workflow from slides to dishes, five immersions into ultrapure water were replaced by one wash with 1 ml ultrapure water. The acquired force-time curves were processed using the JPK SPM data processing software (Bruker, Version 7.0.97), fitting the range of -0.25 µm to 1 µm around the contact point with a Hertz model to yield the Young’s modulus. Indentation curves lacking an identifiable contact point or smooth baseline were excluded from the analysis.

### Dataset registration via image registration

The direct outputs of our two protocols were a 20X Hoechst image with mapped stiffness data acquired during AFM measurements (**Fig. 1c**) and a 10X H&E tile scan of the Visium slide with mapped ST data (**Fig. 1f**). To enable registration of these images, we additionally performed an H&E stain of the tissue slice used for AFM, and imaged it at a Nikon Ti2 inverted microscope with a Plan Apo λ 20X objective (Fig.1d) and as a tilescan with a Plan Apo λ 10X objective (**Fig. 1e**). This allowed us to perform the registration according to the following steps:

1. The 20X Hoechst images with the AFM data (**Fig. 1c**) were mapped to the corresponding 20X H&E images (**Fig. 1d**) of the same regions via affine transformation with the affinder plug-in (v 0.5.0) in napari (v 0.6.6)^71^. Here, nuclei were used as landmarks and a minimum of 10 were selected per image. The plug-in then outputted a transformation matrix in the following format (col = column; trans = translation):

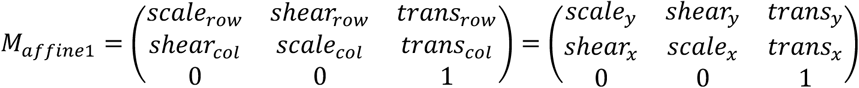
2. The 20X H&E images of the AFM tissue (**Fig. 1d**) were scaled down by a factor 0.5 and mapped to the corresponding 10X H&E tile scan (**Fig. 1e**) via template matching using the TM_CCOEFF_NORMED algorithm in the cv2.matchTemplate module from OpenCV (version 4.10.0.84). The cv2.minMaxLoc function was applied to the result to determine the required translation for the template matching matrix M_tm2_:

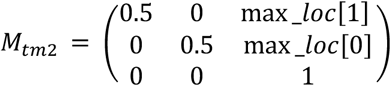
3. The 10X H&E tile scan of the AFM section (**Fig. 1e**) was mapped to the 10X H&E tile scan of the spatial transcriptomic section (**Fig. 1f**) using the napari affinder plug-in. This time, larger structures such as glandular patterns or tumour parenchyma stroma borders were used as landmarks (**Supplementary Fig. 3**). The yielded affine transformation matrix M_affine3_ was in the same format as M_affine1_.

Multiplying the matrices yielded from each transformation step yielded an image transformation matrix:

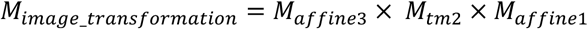

For patient 1, the sample was accidentally rotated by 18° between acquiring images of the H&E-stained AFM-probed region of interest and the H&E-stained full AFM section. Therefore, the image was rotated by 18° for the template matching. To compensate for this, a rotation matrix was multiplied with each yielded template matching matrix M_tm2_. The rotation matrix was computed using the cv2.getRotationMatrix2D function in OpenCV (version 4.10.0.84).

We used the final image transformation matrix with the scipy.ndimage.affine_transform function from the SciPy package (v 1.12.0)^72^ to directly map the 20X Hoechst AFM image to the Visium tilescan. We confirmed proper alignment by visually inspecting an overlay of the affine-transformed 20X Hoechst AFM image with the Visium tilescan.

For each probed FOV, AFM coordinates were generated by creating a grid of coordinates spaced 20 µm apart with its centre in the centre of the AFM Hoechst image. The recorded stiffness values were linked to those coordinates. In order to transform AFM measurement coordinates into the Visium coordinate space, the x and y coordinates of the image transformation matrix needed to be swapped due to the convention of displaying images with the axis order (row, column), i.e. (y,x), but points as (x,y). The resulting matrix M_point_ was then multiplied with each AFM measurement coordinate Point_AFM_space_ to yield the transformed coordinate Point_Visium_space_.

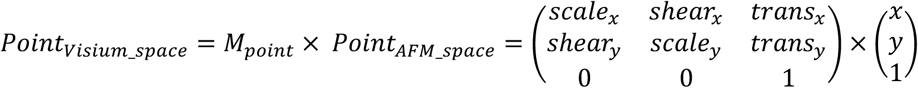

Each transformed AFM coordinate was then mapped to the closest ST spot based on the spot centre coordinates which were acquired as outputs of the SpaceRanger pipeline. Then, error propagation through all transformation steps was performed, accounting for the two affine transformation steps via M_affine1_ and M_affine3_, but assuming no error in the scaling and translation with M_tm2_.

For each affine transformation, a Jacobian matrix J was constructed where each odd row represented the partial derivatives of x’, and each even row represented the partial derivatives of y’. x and y were the original landmark coordinates and x’ and y’ the affine transformed ones.

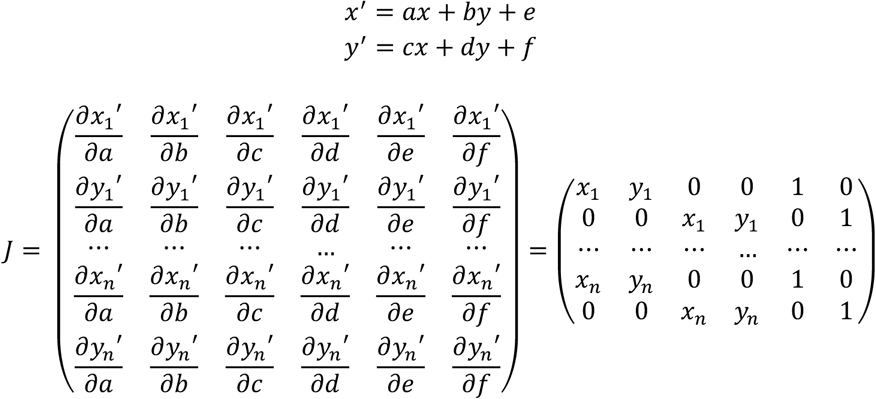

From the Jacobian matrix, the landmark residuals r and the degrees of freedom (2n-6), the covariance matrix Σ_p_ was computed. σ^2^ refers to the estimated variance of the landmarks.

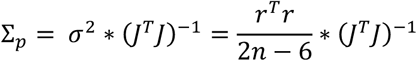

Having constructed all covariance matrices, we performed an error propagation. For a point P_A_ that represents an AFM coordinate that is then affine transformed into point P_B_, transformed with the matrix acquired from template matching to point P_C_, and affine transformed again to the final coordinate in the Visium space to P_D_,, the standard deviations σ_x_ and σ_y_ were extracted from the propagated covariance matrix Σ_pD_ at point P_D_. Σ_p1_ and Σ_p2_ are the corresponding parameter covariance matrices, and J_PA_ and J_PC_ represent the Jacobian Matrices for Point P_A_ and P_C_, respectively. From the transformation matrices M_affine1_, M_tm2_ and M_affine3_, only the scale and shear components M^ss^ were considered.

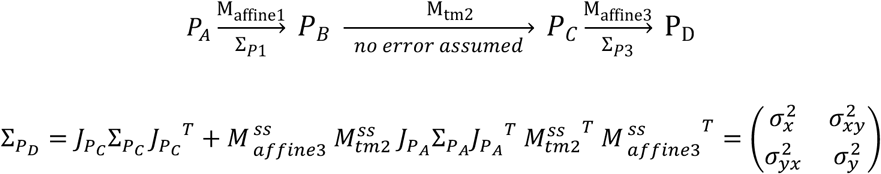

To refine our dataset, we chose a mapping error of 0.5 σ, meaning that we only kept coordinates that can be shifted by ±0.5 σ_x_, ±0.5 σ_y_ or both, and after each shift were still closest to the same ST spot. 3521 out of 4584 mapped measurements were kept. With our method, measurements that were mapped between the physical ST spots were still included as long as their mapping error was small enough to still consistently map them to the same spot when moving the measurements by its mapping errors.

AFM measurements were grouped based on corresponding Visium capture spots. Groups with only two measurements were retained if their coefficient of variation was below 40%. For groups with three or more measurements, outliers exceeding 3.5 median absolute deviations (MADs) from the group median and the two spots exceeding a mean stiffness of 12500 Pa were removed. The mean value of the measurements was then calculated for each group and linked to their corresponding ST spot.

### Differential gene expression analysis

As established previously, conventional differential expression analysis tools tend to underperform when applied to zero-inflated single-cell expression data with a high dropout rate^73^. Therefore, for our ST dataset, we opted to first infer gene- and cell-specific weights using the ZINB-WaVE method. In detail, after the removal of mitochondrial and ribosomal genes, one thousand highly variable genes were identified by the method and used to calculate observational weights with ZINB-WaVE. These weights were next incorporated into edgeR along with raw counts to calculate differentially expressed genes using a log fold change threshold of 0.5 and FDR threshold of 5%. The comparison was done across all 11 samples combined between “high stiffness” and “low stiffness” groups of capture spots based on three thresholds of 260 Pa (the rounded median stiffness of our 297 spot curated dataset), 520 Pa and 780 Pa. We additionally performed this analysis on datasets with reduced mapping radii (27.5 µm and 38.75 µm) using an FDR threshold of 10% and 5%, respectively.

### Random Forest regression

The dataset was divided into features (gene expression) and the target variable (mean stiffness value). A RandomForestRegressor model from scikit-learn (v1.5.1) was employed, with 100 estimators and a maximum depth of 10. A 5-fold cross-validation was performed to evaluate the model’s performance, calculating both the Root Mean Squared Error (RMSE) and R-squared values for each fold and on average. After cross-validation, the model was fitted to the entire dataset to identify and rank the importance of each gene in predicting the stiffness. The partial dependence of the top ten most important genes was estimated.

### Statistical analysis

All measurements were taken from distinct samples: 11 CRC LM tissue samples from 10 patients (with Patient 7 contributing two samples from peripheral and central regions of the same metastasis). For all statistical analysis a p-value < 0.05 was considered statistically significant.

On the data displayed in **Fig. 2d**, statistical analyses were performed using R (version 4.3.1). First, a one-way ANOVA was performed to assess differences in mean stiffness between ROIs, with ROI as the explanatory variable. The residual variation was used to estimate the variability within ROIs, representing differences between sections. From these results we computed the standard deviation displayed in **Fig. 3e**. To assess a potential systematic contribution of the section to variations in stiffness, a two-way ANOVA using the ROI and section as factors was subsequently performed.

For the Boxplot in **Fig. 3g**, statistical analyses were performed using Python (version 3.13.12). Normality of data distributions was assessed using the Shapiro-Wilk test, and homogeneity of variances was assessed using Levene’s test. As these assumptions were not met, comparisons were performed using a non-parametric Kruskal-Wallis test. The lower and upper hinges correspond to the first and third quartiles (the 25th and 75th percentiles); the whiskers extend to the largest and smallest values no further than 1.5× the interquartile range (IQR) from the hinge.

Linear regressions were performed on log-transformed data (**Fig. 4b-I**, **Fig. 5g,i-l**) or normalized data (**Supplementary Fig. 1**) using the lineregress function from the SciPy package in Python (version 3.13.12). The correlation strength was assessed via the Pearsons’s correlation coefficient r.

## Code availability

All custom code required for the mapping, error propagation, differential gene expression analysis and random forest regression is available in the MechanoMaST GitHub repository (https://github.com/diz-lab/MechanoMaST). The version used for this manuscript (v.1.0.0) is archived on Zenodo (doi.org/<u>10.5281/zenodo.23064485</u>).

## Data availability

Raw and processed Visium data as well as full-resolution images of the tissues will be deposited to a database upon formal publication of this manuscript.

## Supplementary Information

**Supplementary Figure 1:**
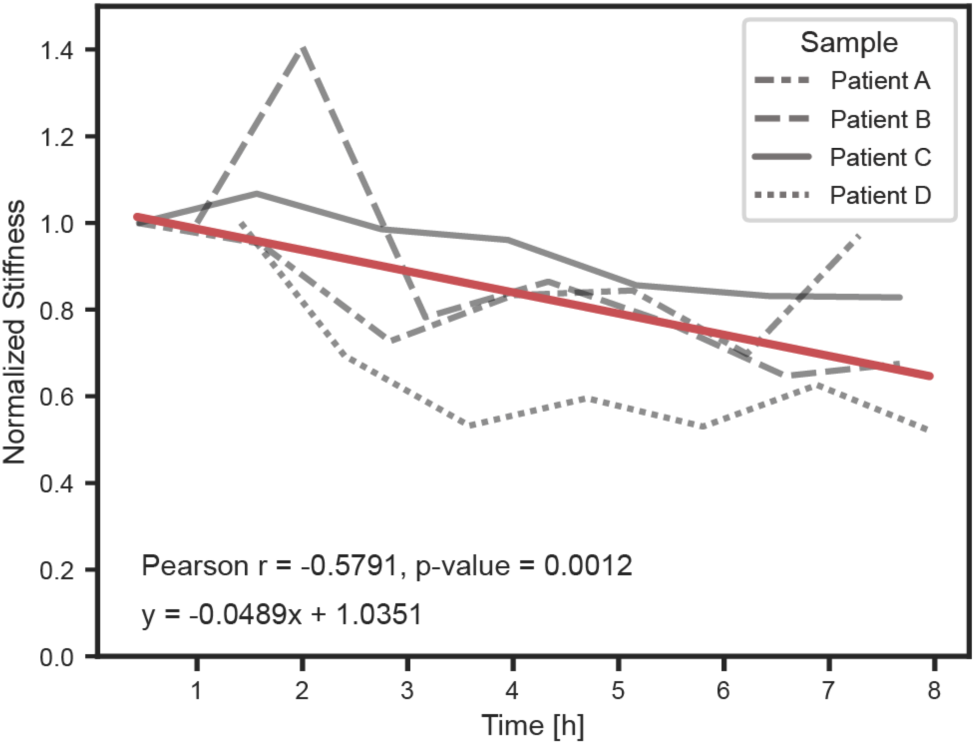
Changes of tissue stiffness over time. The stiffness values shown in Figure 2f were normalized per patient and a linear regression was performed. The time refers to the time after thawing the tissue.

**Supplementary Figure 2:**
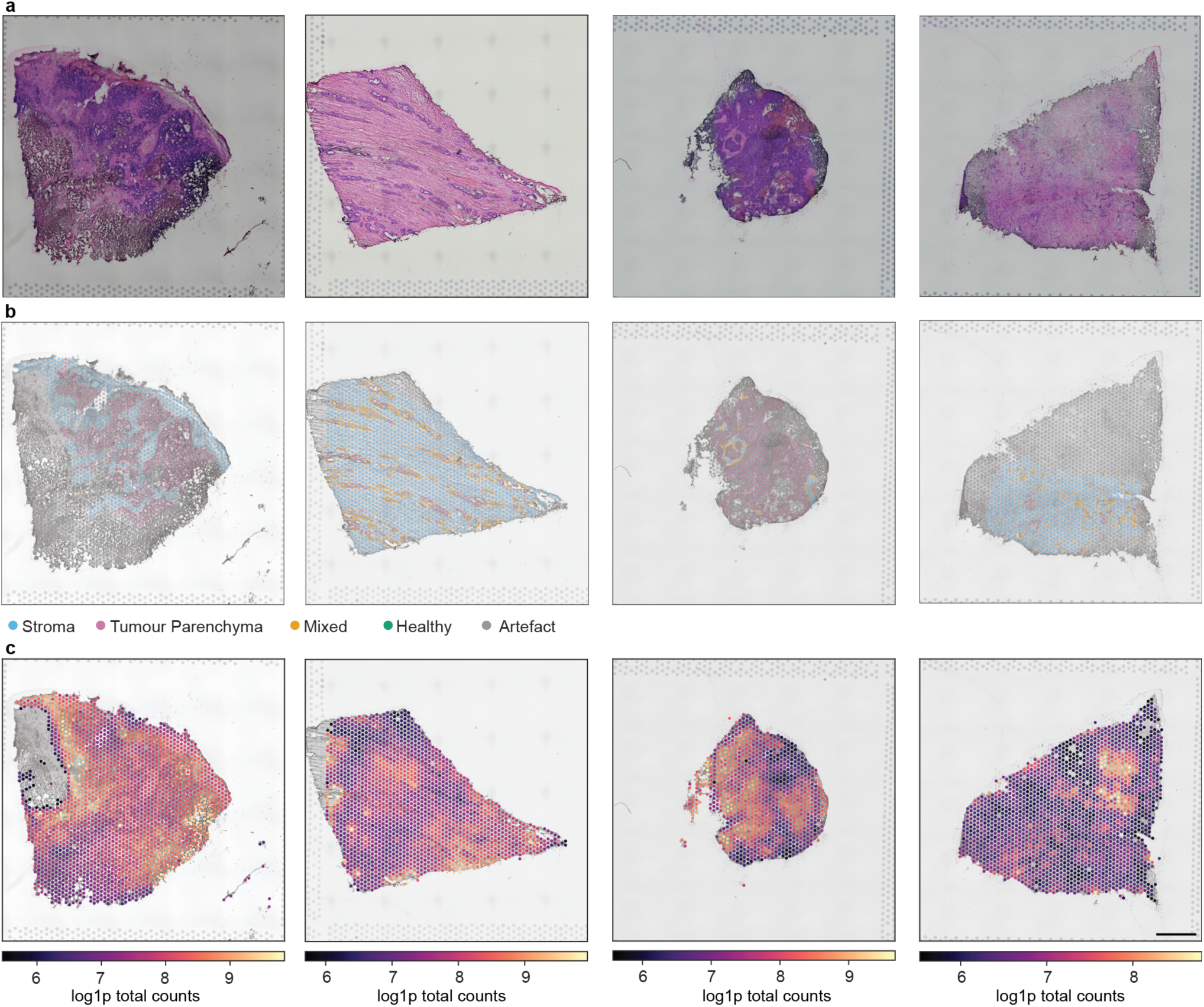
Spatial heterogeneity of CRC LM samples across patients. (4 of 11 CRC LM samples shown: samples 1, 7b, 9 and 10, from left to right; full QC metrics for all 11 samples are provided in **Supplementary Table 2**, total number of spots: 15890). **a**, 20 µm CRC LM cryosections on Visium chips stained with H&E. **b**, Pathologist’s annotations of samples. The bottom and left of sample 1 were labelled as artefacts due to cryodamage. The top and left of sample 10 were labelled as artefacts due to edema. **c**, Log1p of total counts. The top left of sample 1 was excluded due to improper permeabilisation. Scale bar: 1 mm.

**Supplementary Figure 3:**
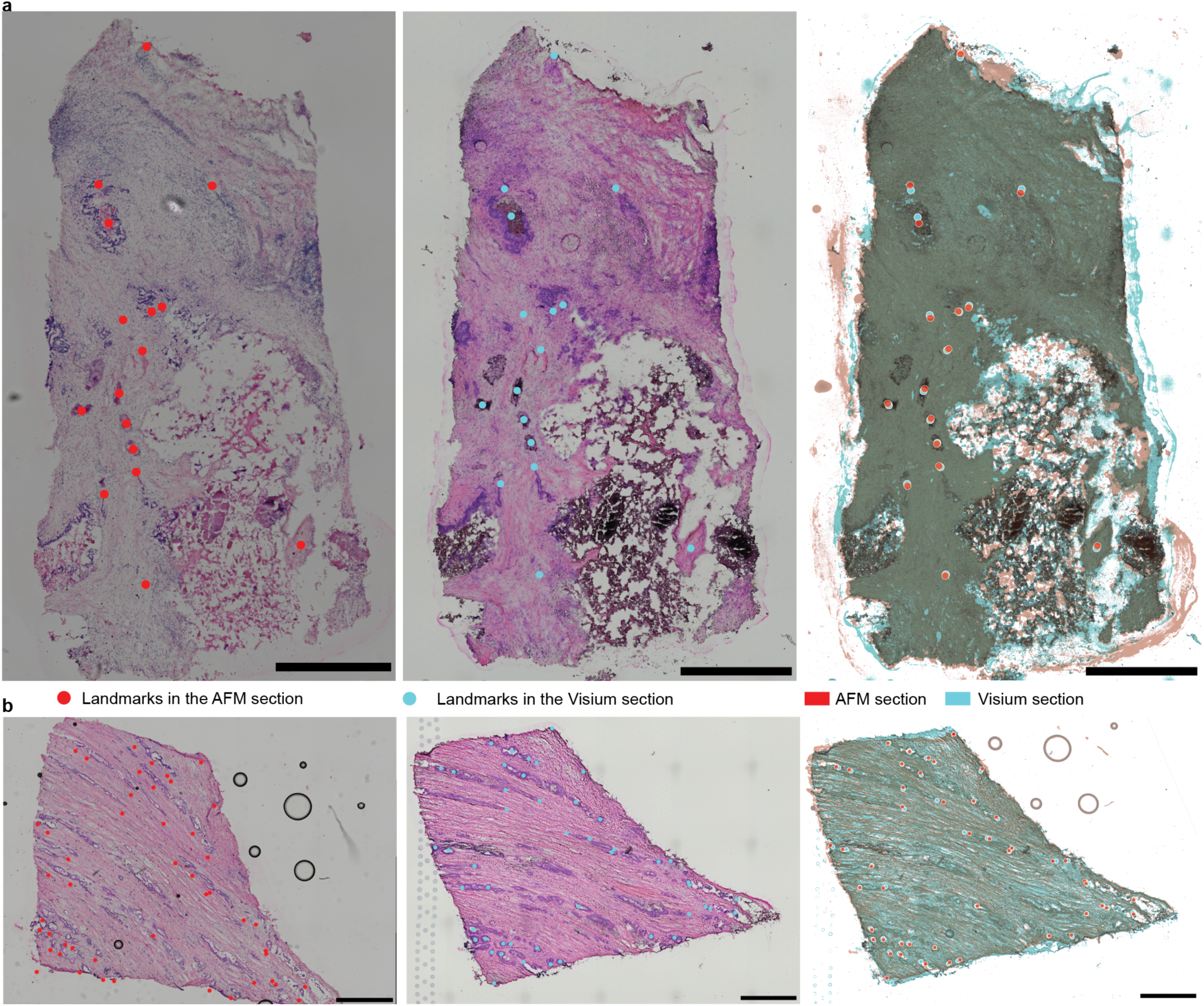
Landmark-based affine transformations of directly adjacent sections. (2 representative examples of the 11 CRC LM sample pairs processed with MechanoMaST; see **Methods** for the full registration pipeline applied to all samples). **a**,**b**, Directly adjacent CRC LM cryosection of Patient 5 (a) and 7b (b). The individual sections including manually selected landmarks as well as an overlay of the sections is shown. Scale bars: 1 mm.

**Supplementary Figure 4:**
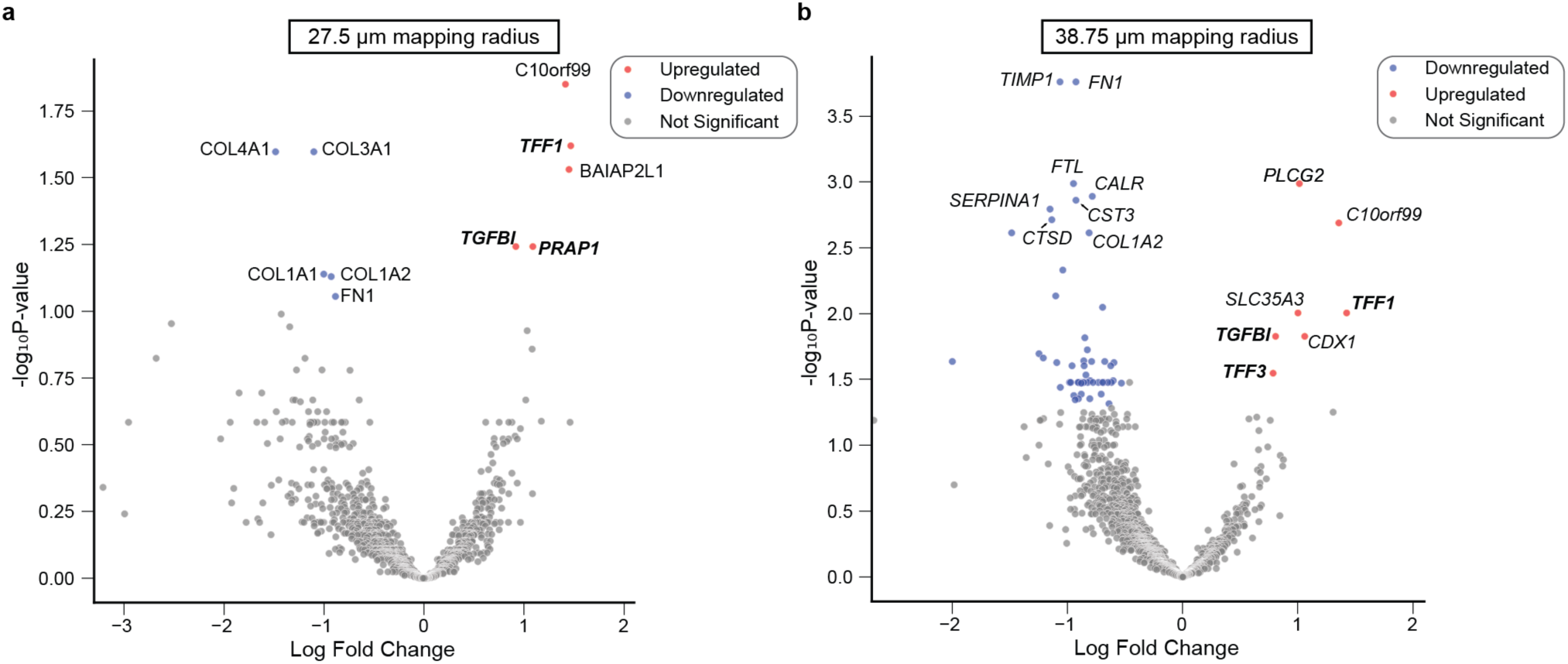
Verification of candidates using different mapping radii. (n = 228 ST spots or 273 ST spots for the 27.5 µm and the 38.75 µm mapping radius, respectively)**. a**,**b,** Differential gene expression analysis. Spots above 260 Pa were considered as stiff. Red and blue dots refer to genes that are upregulated or downregulated, respectively, in stiff spots.

**Supplementary Figure 5:**
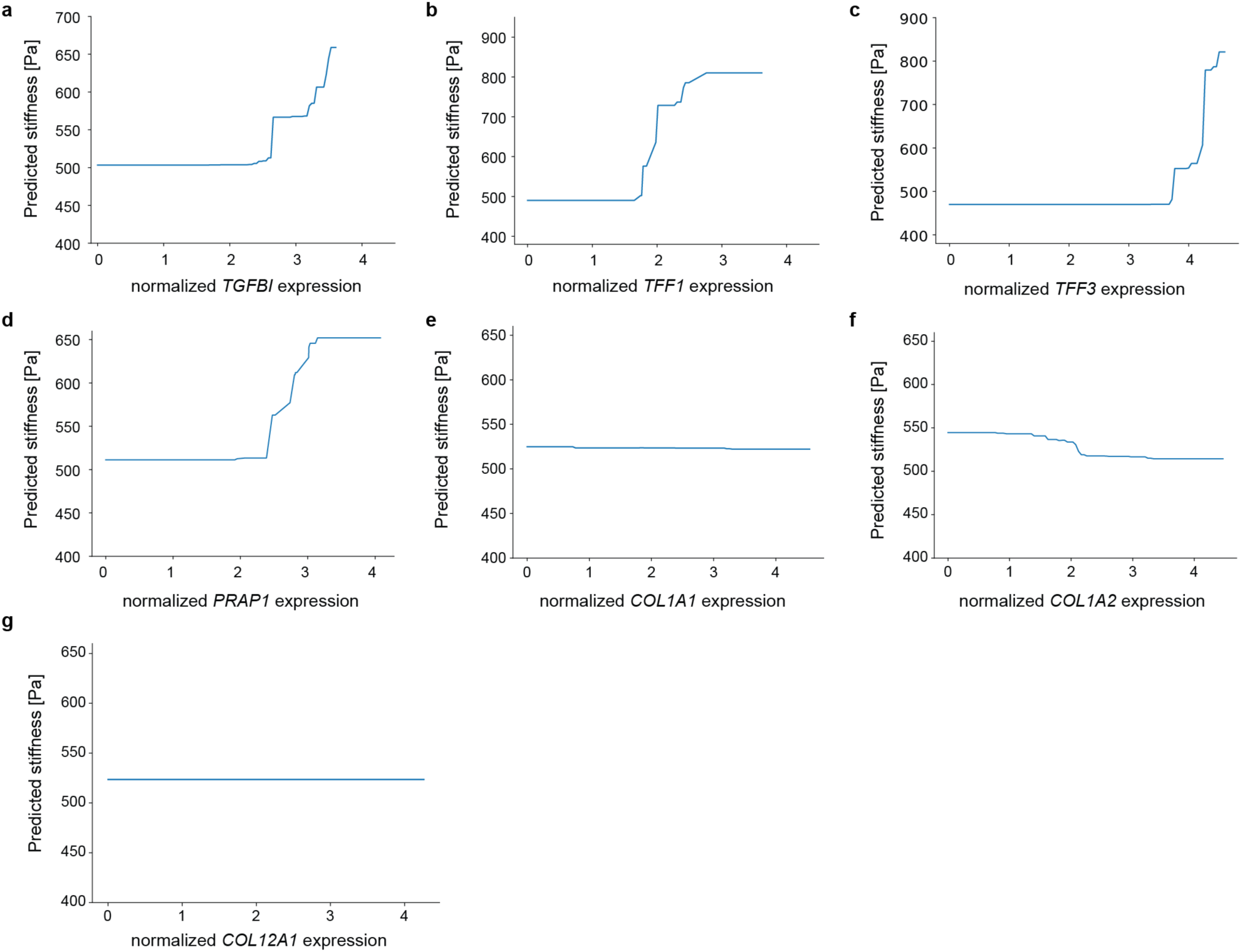
Stiffness predicted by the random forest model in response to the lognorm expression level of the candidate genes or other ECM components. (n = 297 ST spots per panel, matching the dataset used for the linear regressions in Fig. 4b**–h**). **a**, Candidate gene *TGFBI;* **b**, *TFF1;* **c**, *TFF3*; **d**, *PRAP1*; **e**, ECM component *COL1A1*; **f,** ECM component *COL1A2*; **g**, ECM component *COL12A1*.

**Supplementary Figure 6:**
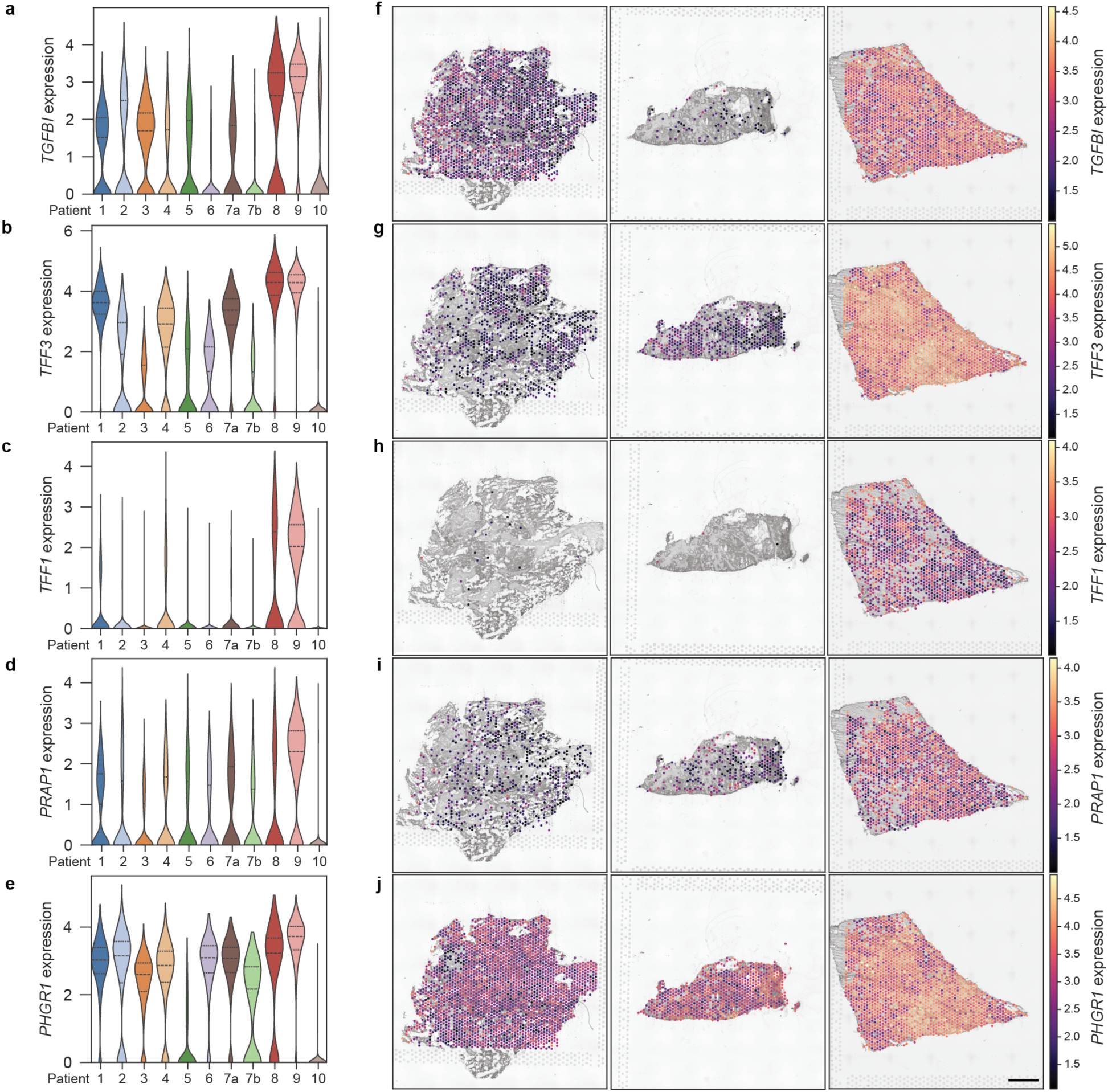
Candidate gene expression levels across patients. (n = 11 CRC LM samples from 10 patients, matching the full MechanoMaST dataset, including non-stromal spots). **a-e**, Expression of target genes across patients. *TGFBI*, *TFF3*, *PRAP1*, and *PHGR1* show expression across patients, indicating that their potential stiffness contribution is not patient-specific. *TFF1* is predominantly expressed in Patient 7, which could suggest a patient-specific role. **f-j** Representative spatial expression maps for three samples (left column: Patient 3; middle column: Patient 7b; bottom column: Patient 8). Scale bar: 1mm.

**Supplementary Table 1:** Patient characteristics.

| Sex | Male | Female |
| --- | --- | --- |
|  | 8 | 2 |
| Age [years] | < 60 | > 60 |
|  | 5 | 5 |
| T-Stage of the primary<br>tumour (2 missing) | T2 | T3 |
|  | 3 | 5 |
| Localization of the primary<br>tumour | Colon | Rectum |
|  | 6 | 4 |

**Supplementary Table 2:**
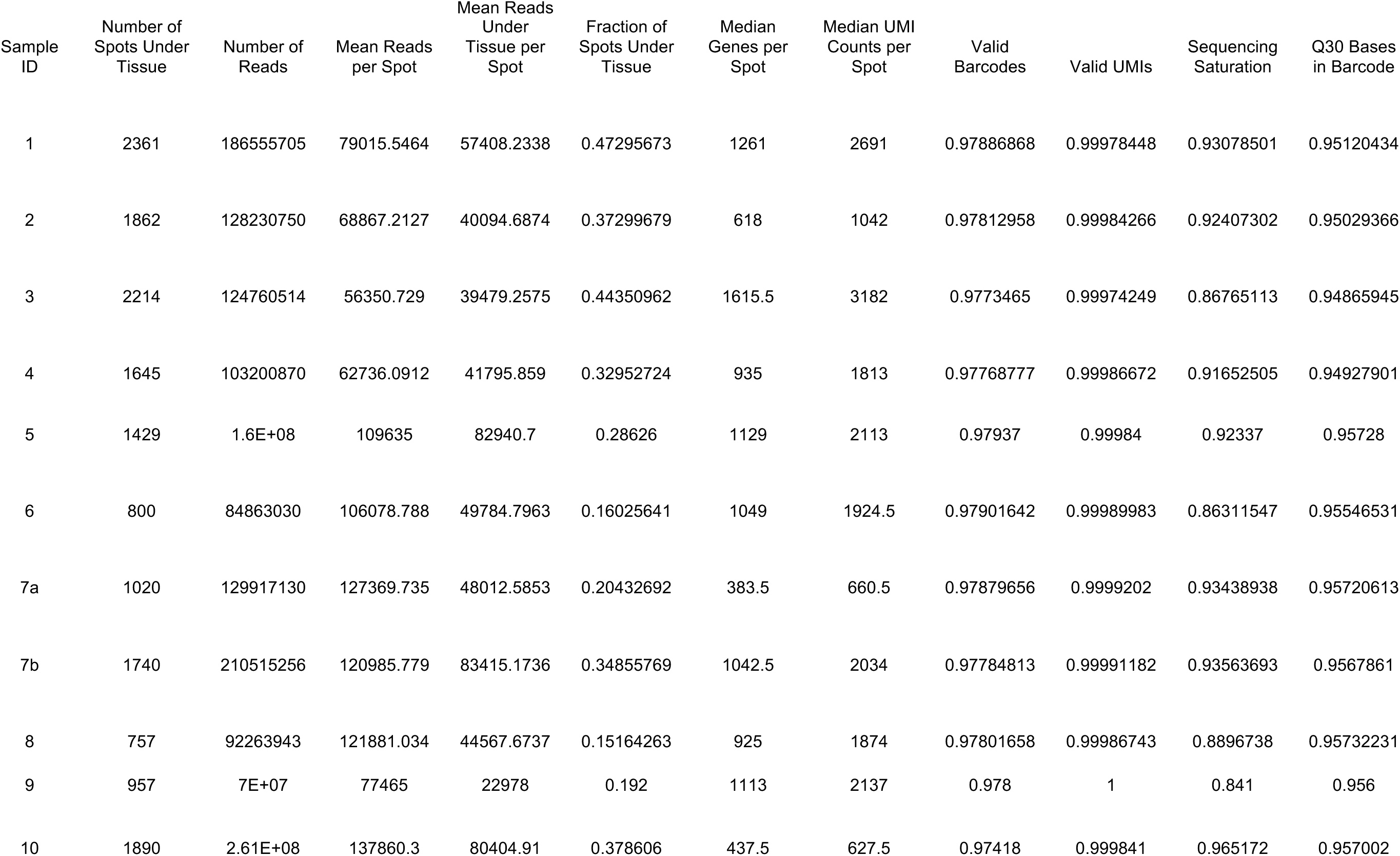
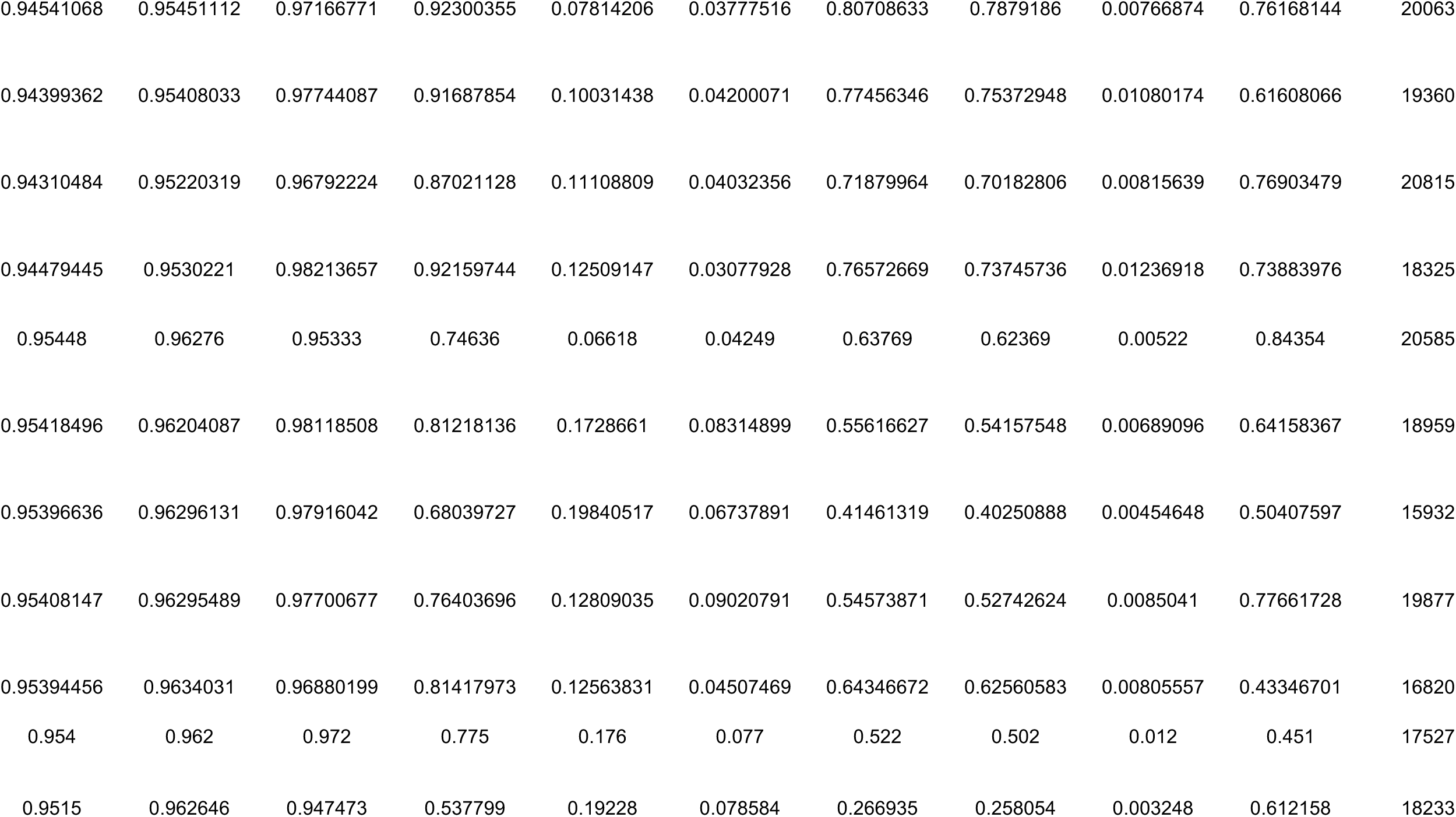
Visium quality control parameters.

**Supplementary Table 3:** AFM quality control parameters. The mapping error refers to half the standard deviation from the error propagation. In total, 5810 AFM measurements were conducted, 4584 were successfully fitted, and 3521 were successfully mapped with a small enough error.

| Sample ID | ROI | Conducted Measurements | Fitted Measurements | Average RMS [pN] | Mapped Measurements | Average Mapping error [µm] |
| --- | --- | --- | --- | --- | --- | --- |
| 1 | ROI 1 | 100 | 80 | 16.01951 | 63 | 10.26554 |
|  | ROI 2 | 100 | 82 | 13.64194 | 51 | 14.813315 |
|  | ROI 3 | 100 | 74 | 10.51639 | 36 | 20.46147 |
|  | ROI 4 | 68* | 43 | 15.77103 | 38 | 7.90978 |
|  | ROI 5 | 225 | 175 | 11.30792 | 150 | 7.751055 |
| 2 | ROI 1 | 100 | 68 | 17.69785 | 33 | 20.219775 |
|  | ROI 2 | 100 | 82 | 15.18711 | 69 | 7.669525 |
|  | ROI 3 | 100 | 84 | 12.65432 | 16 | 33.898555 |
|  | ROI 4 | 100 | 77 | 14.06616 | 64 | 8.632645 |
| 3 | ROI 1 | 100 | 82 | 14.91092 | 69 | 6.432905 |
|  | ROI 2 | 100 | 82 | 10.78847 | 75 | 5.05822 |
|  | ROI 3 | 100 | 85 | 12.54754 | 61 | 13.29735 |
|  | ROI 4 | 100 | 82 | 11.76794 | 68 | 9.75022 |
|  | ROI 5 | 100 | 71 | 12.77581 | 67 | 5.939535 |
|  | ROI 6 | 25 | 19 | 10.7979 | 18 | 5.729555 |
| 4 | ROI 1 | 100 | 78 | 14.95035 | 74 | 5.564175 |
|  | ROI 2 | 100 | 78 | 13.14165 | 74 | 5.681335 |
|  | ROI 3 | 100 | 83 | 13.05696 | 61 | 10.985165 |
|  | ROI 4 | 100 | 76 | 9.89384 | 73 | 2.945122 |
| 5 | ROI 1 | 100 | 81 | 8.601266 | 73 | 6.93034 |
|  | ROI 2 | 100 | 84 | 10.28374 | 69 | 7.735965 |
|  | ROI 3 | 100 | 87 | 9.060281 | 43 | 19.424065 |
|  | ROI 4 | 100 | 74 | 10.03925 | 20 | 31.22907 |
|  | ROI 5 | 100 | 75 | 9.441483 | 39 | 21.47206 |
|  | ROI 6 | 100 | 72 | 9.756373 | 25 | 26.017285 |
|  | ROI 7 | 100 | 83 | 10.72353 | 75 | 6.6559 |
| 6 | ROI 1 | 100 | 87 | 13.51705 | 61 | 12.289295 |
|  | ROI 2 | 100 | 80 | 13.79854 | 62 | 10.839205 |
|  | ROI 3 | 100 | 84 | 11.5631 | 67 | 8.79578 |
|  | ROI 4 | 100 | 77 | 19.17242 | 73 | 4.6085915 |
|  | ROI 5 | 100 | 79 | 16.33614 | 73 | 6.860575 |
|  | ROI 6 | 100 | 85 | 12.42707 | 65 | 12.547545 |
| 7a | ROI 1 | 400 | 321 | 13.80221 | 291 | 6.175565 |
|  | ROI 2 | 100 | 79 | 12.00101 | 65 | 8.869725 |
| 7b | ROI 1 | 100 | 81 | 35.06147 | 70 | 8.375475 |
|  | ROI 2 | 100 | 82 | 20.90996 | 70 | 7.22708 |
|  | ROI 3 | 100 | 76 | 28.31479 | 69 | 4.9020155 |
|  | ROI 4 | 100 | 78 | 14.77609 | 69 | 6.843845 |
|  | ROI 5 | 100 | 76 | 14.1347 | 67 | 6.19456 |
|  | ROI 6 | 100 | 78 | 18.36346 | 75 | 3.6971645 |
|  | ROI 7 | 100 | 69 | 26.8894 | 67 | 4.278782 |
| 8 | ROI 1 | 200 | 162 | 15.23238 | 88 | 16.29735 |
|  | ROI 2 | 100 | 83 | 13.48017 | 75 | 4.8752275 |
|  | ROI 3 | 162 | 127 | 14.54107 | 115 | 5.91306 |
|  | ROI 4 | 25 | 18 | 12.77409 | 15 | 15.30023 |
|  | ROI 5 | 50 | 35 | 15.82451 | 34 | 3.8778025 |
| 9 | ROI 1 | 64 | 53 | 15.7878 | 52 | 4.1520945 |
|  | ROI 2 | 25 | 22 | 27.05802 | 22 | 3.566545 |
|  | ROI 3 | 49 | 37 | 17.05474 | 27 | 9.651365 |
|  | ROI 4 | 49 | 40 | 14.67031 | 31 | 10.22222 |
|  | ROI 5 | 36 | 31 | 11.44762 | 29 | 5.09209 |
|  | ROI 6 | 16 | 16 | 11.65008 | 15 | 5.94029 |
|  | ROI 7 | 16 | 15 | 17.66657 | 14 | 4.6640405 |
| 10 | ROI 1 | 100 | 80 | 17.83046 | 46 | 18.593135 |
|  | ROI 2 | 100 | 81 | 16.07811 | 46 | 15.90781 |
|  | ROI 3 | 100 | 74 | 17.6199 | 33 | 20.573785 |
|  | ROI 4 | 100 | 79 | 17.72498 | 52 | 13.49829 |
|  | ROI 5 | 100 | 64 | 18.97355 | 41 | 17.452655 |
|  | ROI 6 | 100 | 78 | 18.03532 | 38 | 17.14624 |

\* 100 measurements were originally conducted, but 32 were excluded due to being located in an area that was excluded from the dataset due to improper tissue permeabilization during the Visium workflow.

**Supplementary Tables 4-9 can be found in an additional excel file.**

## References

1. Barriga, E. H., Franze, K., Charras, G. & Mayor, R. Tissue stiffening coordinates morphogenesis by triggering collective cell migration in vivo. Nature 554, 523–527 (2018).

2. Shellard, A. & Mayor, R. Collective durotaxis along a self-generated stiffness gradient in vivo. Nature 600, 690–694 (2021).

3. Mitrossilis, D. et al. Mechanotransductive cascade of Myo-II-dependent mesoderm and endoderm invaginations in embryo gastrulation. Nat Commun 8, 13883 (2017).

4. Engler, A. J., Sen, S., Sweeney, H. L. & Discher, D. E. Matrix Elasticity Directs Stem Cell Lineage Specification. Cell 126, 677–689 (2006).

5. Liang, R. & Song, G. Matrix stiffness-driven cancer progression and the targeted therapeutic strategy. Mechanobiol Med 1, 100013 (2023).

6. Santos, A. & Lagares, D. Matrix Stiffness: the Conductor of Organ Fibrosis. Curr Rheumatol Rep 20, 2 (2018).

7. Al-Hilal, T. A. et al. Durotaxis is a driver and potential therapeutic target in lung fibrosis and metastatic pancreatic cancer. Nat Cell Biol 27, 1543–1554 (2025).

8. Pavuluri, K. et al. Brain Mechanical Properties Predict Longitudinal Cognitive Change in Aging and Alzheimer’s Disease. Neurobiol Aging 147, 203–212 (2025).

9. Donnaloja, F. et al. Unravelling the mechanotransduction pathways in Alzheimer’s disease. J Biol Eng 17, 22 (2023).

10. Method of the Year 2019: Single-cell multimodal omics. Nat Methods 17, 1–1 (2020).

11. Method of the Year 2020: spatially resolved transcriptomics. Nat Methods 18, 1–1 (2021).

12. Lee, D. et al. Multi-omics single-cell analysis reveals key regulators of HIV-1 persistence and aberrant host immune responses in early infection. eLife 14, RP104856 (2025).

13. Xie, Y. et al. Noninvasive prognostic classification of ITH in HCC with multi-omics insights and therapeutic implications. Sci. Adv. 11, eads8323 (2025).

14. Wang, X. et al. Single-cell multi-omics sequencing uncovers region-specific plasticity of glioblastoma for complementary therapeutic targeting. Sci. Adv. 10, eadn4306 (2024).

15. Shi, Z. et al. Single-nucleus multi-omics identifies shared and distinct pathways in Pick’s and Alzheimer’s disease. Sci. Adv. 11, eads7973 (2025).

16. Heide, T. et al. The co-evolution of the genome and epigenome in colorectal cancer. Nature 611, 733–743 (2022).

17. Daucke, R. et al. Refining spatial proteomics by mass spectrometry: an efficient workflow tailored for archival tissue. BMC Methods 3, 20 (2026).

18. Ståhl, P. L. et al. Visualization and analysis of gene expression in tissue sections by spatial transcriptomics. Science 353, 78–82 (2016).

19. Hallou, A., He, R., Simons, B. D. & Dumitrascu, B. A computational pipeline for spatial mechano-transcriptomics. Nat Methods 22, 737–750 (2025).

20. Mathavan, N. et al. Spatial transcriptomics in bone mechanomics: Exploring the mechanoregulation of fracture healing in the era of spatial omics. Sci. Adv. 11, eadp8496 (2025).

21. Stashko, C. et al. A convolutional neural network STIFMap reveals associations between stromal stiffness and EMT in breast cancer. Nat Commun 14, 3561 (2023).

22. Krieg, M. et al. Atomic force microscopy-based mechanobiology. Nat Rev Phys 1, 41–57 (2018).

23. Kawano, S. et al. Assessment of elasticity of colorectal cancer tissue, clinical utility, pathological and phenotypical relevance. Cancer Sci 106, 1232–1239 (2015).

24. Shen, Y. et al. Reduction of Liver Metastasis Stiffness Improves Response to Bevacizumab in Metastatic Colorectal Cancer. Cancer Cell 37, 800–817.e7 (2020).

25. Loroña, N. C., Sankar, K., Stern, M. C., Schmit, S. L. & Figueiredo, J. C. de novo metastases in patients with primary colorectal cancer: a Surveillance, Epidemiology, and End Results analysis. Cancer Causes Control 36, 937–946 (2025).

26. Cañellas-Socias, A., Sancho, E. & Batlle, E. Mechanisms of metastatic colorectal cancer. Nat Rev Gastroenterol Hepatol 21, 609–625 (2024).

27. Deville, S. S. & Cordes, N. The Extracellular, Cellular, and Nuclear Stiffness, a Trinity in the Cancer Resistome— A Review. Front Oncol 9, 1376 (2019).

28. Van Huizen, N. A. et al. Up-regulation of collagen proteins in colorectal liver metastasis compared with normal liver tissue. Journal of Biological Chemistry 294, 281–289 (2019).

29. Kalli, M., Poskus, M. D., Stylianopoulos, T. & Zervantonakis, I. K. Beyond matrix stiffness: targeting force-induced cancer drug resistance. Trends in Cancer 9, 937–954 (2023).

30. Gamradt, P., et al. Stiffness-induced cancer-associated fibroblasts are responsible for immunosuppression in a platelet-derived growth factor ligand-dependent manner. PNAS Nexus 2, pgad405 (2023).

31. Mai, Z., Lin, Y., Lin, P., Zhao, X. & Cui, L. Modulating extracellular matrix stiffness: a strategic approach to boost cancer immunotherapy. Cell Death Dis 15, 307 (2024).

32. Hastings, J. F., Skhinas, J. N., Fey, D., Croucher, D. R. & Cox, T. R. The extracellular matrix as a key regulator of intracellular signalling networks. Br J Pharmacol 176, 82–92 (2019).

33. Feng, X. et al. Targeting extracellular matrix stiffness for cancer therapy. Front Immunol 15, 1467602 (2024).

34. Zhang, M. & Zhang, B. Extracellular matrix stiffness: mechanisms in tumor progression and therapeutic potential in cancer. Exp Hematol Oncol 14, 54 (2025).

35. Özdemir, B. C. et al. Depletion of Carcinoma-Associated Fibroblasts and Fibrosis Induces Immunosuppression and Accelerates Pancreas Cancer with Diminished Survival. Cancer Cell 25, 719–734 (2014).

36. Rhim, A. D. et al. Stromal Elements Act to Restrain, Rather Than Support, Pancreatic Ductal Adenocarcinoma. Cancer Cell 25, 735–747 (2014).

37. Krishnamurty, A. T. et al. LRRC15+ myofibroblasts dictate the stromal setpoint to suppress tumour immunity. Nature 611, 148–154 (2022).

38. Yan, W. et al. CAFs activated by YAP1 upregulate cancer matrix stiffness to mediate hepatocellular carcinoma progression. J Transl Med 23, 450 (2025).

39. Martinez-Vidal, L. et al. Causal contributors to tissue stiffness and clinical relevance in urology. Commun Biol 4, 1011 (2021).

40. Pavuluri, K. et al. Differential effect of dementia etiology on cortical stiffness as assessed by MR elastography. NeuroImage: Clinical 37, 103328 (2023).

41. Lebrigand, K. et al. The spatial landscape of gene expression isoforms in tissue sections. Nucleic Acids Res 51, e47 (2023).

42. Ning, Y. et al. Gastrointestinal pan-cancer landscape of tumor matrix heterogeneity identifies biologically distinct matrix stiffness subtypes predicting prognosis and chemotherapy efficacy. Computational and Structural Biotechnology Journal 21, 2744–2758 (2023).

43. Shen, Y. et al. Matrix stiffness-related extracellular matrix signatures and the DYNLL1 protein promote hepatocellular carcinoma progression through the Wnt/β-catenin pathway. BMC Cancer 24, 1211 (2024).

44. Goehrig, D. et al. Stromal protein βig-h3 reprogrammes tumour microenvironment in pancreatic cancer. Gut 68, 693–707 (2019).

45. Chen, J. G. et al. Giotto Suite: a multiscale and technology-agnostic spatial multiomics analysis ecosystem. Nat Methods 22, 2052–2064 (2025).

46. Marconato, L. et al. SpatialData: an open and universal data framework for spatial omics. Nat Methods 22, 58–62 (2025).

47. Ravi, V. M. et al. Spatially resolved multi-omics deciphers bidirectional tumor-host interdependence in glioblastoma. Cancer Cell 40, 639–655.e13 (2022).

48. Lotz, J., Weiss, N., van der Laak, J. & Heldmann, S. Comparison of consecutive and restained sections for image registration in histopathology. J Med Imaging (Bellingham) 10, 067501 (2023).

49. Oliveira, M. F. de et al. High-definition spatial transcriptomic profiling of immune cell populations in colorectal cancer. Nat Genet 57, 1512–1523 (2025).

50. Shioka, I., et al. *Ex vivo* SIM-AFM measurements reveal the spatial correlation of stiffness and molecular distributions in 3D living tissue. Acta Biomaterialia 189, 351–365 (2024).

51. Plodinec, M. et al. The nanomechanical signature of breast cancer. Nature Nanotech 7, 757–765 (2012).

52. Calò, A. et al. Spatial mapping of the collagen distribution in human and mouse tissues by force volume atomic force microscopy. Sci Rep 10, 15664 (2020).

53. Verzijl, N. et al. Effect of Collagen Turnover on the Accumulation of Advanced Glycation End Products*. Journal of Biological Chemistry 275, 39027–39031 (2000).

54. Buccitelli, C. & Selbach, M. mRNAs, proteins and the emerging principles of gene expression control. Nat Rev Genet 21, 630–644 (2020).

55. Ponomarenko, E. A. et al. Workability of mRNA Sequencing for Predicting Protein Abundance. Genes (Basel) 14, 2065 (2023).

56. Prabahar, A., et al. Unraveling the complex relationship between mRNA and protein abundances: a machine learning-based approach for imputing protein levels from RNA-seq data. NAR Genom Bioinform 6, lqae019 (2024).

57. Barallobre-Barreiro, J. et al. Proteomics Analysis of Cardiac Extracellular Matrix Remodeling in a Porcine Model of Ischemia/Reperfusion Injury. Circulation 125, 789–802 (2012).

58. Watson, A. W. et al. Breast tumor stiffness instructs bone metastasis via maintenance of mechanical conditioning. Cell Rep 35, 109293 (2021).

59. Patry, M. et al. βig-h3 Represses T-Cell Activation in Type 1 Diabetes. Diabetes 64, 4212–4219 (2015).

60. Bastholm, S. K. et al. Trefoil factor peptide 3 is positively correlated with the viscoelastic properties of the cervical mucus plug. Acta Obstetricia et Gynecologica Scandinavica 96, 47–52 (2017).

61. LeBaron, R. G. et al. βIG-H3, a Novel Secretory Protein Inducible by Transforming Growth Factor-β, Is Present in Normal Skin and Promotes the Adhesion and Spreading of Dermal Fibroblasts In Vitro. Journal of Investigative Dermatology 104, 844–849 (1995).

62. Zhang, J. et al. The Proline-Rich Acidic Protein Is Epigenetically Regulated and Inhibits Growth of Cancer Cell Lines. Cancer Res 63, 6658–6665 (2003).

63. Braga Emidio, N., Brierley, S. M., Schroeder, C. I. & Muttenthaler, M. Structure, Function, and Therapeutic Potential of the Trefoil Factor Family in the Gastrointestinal Tract. ACS Pharmacol Transl Sci 3, 583–597 (2020).

64. Lee, S. G. et al. TGFBI remodels adipose metabolism by regulating the Notch-1 signaling pathway. Exp Mol Med 55, 520–531 (2023).

65. Bazina, I. et al. The Effect of Tff3 Deficiency on the Liver of Mice Exposed to a High-Fat Diet. Biomedicines 13, 1024 (2025).

66. Šešelja, K. et al. Tff3 Deficiency Protects against Hepatic Fat Accumulation after Prolonged High-Fat Diet. Life (Basel) 12, 1288 (2022).

67. Peng, H. et al. PRAP1 is a novel lipid-binding protein that promotes lipid absorption by facilitating MTTP-mediated lipid transport. J Biol Chem 296, 100052 (2020).

68. Kumar, R. et al. Liver Stiffness Measurements in Patients with Different Stages of Nonalcoholic Fatty Liver Disease: Diagnostic Performance and Clinicopathological Correlation. Dig Dis Sci 58, 265–274 (2013).

69. Wang, Z. et al. Extracellular Vesicles in Fatty Liver Promote a Metastatic Tumor Microenvironment. Cell Metab 35, 1209–1226.e13 (2023).

70. Shen, Y., Schmidt, T. & Diz-Muñoz, A. Protocol on Tissue Preparation and Measurement of Tumor Stiffness in Primary and Metastatic Colorectal Cancer Samples with an Atomic Force Microscope. STAR Protocols 1, 100167 (2020).

71. Sofroniew, N., et al. napari: a multi-dimensional image viewer for Python. Zenodo 10.5281/ZENODO.3555620 (2025).

72. Virtanen, P. et al. SciPy 1.0: fundamental algorithms for scientific computing in Python. Nat Methods 17, 261–272 (2020).

73. Van den Berge, K., et al. Observation weights unlock bulk RNA-seq tools for zero inflation and single-cell applications. Genome Biology 19, 24 (2018).

